# aaRSID, an engineered pyrrolysyl-tRNA synthetase platform for multi-probe proximity proteomics

**DOI:** 10.64898/2026.09.22.753581

**Authors:** Zhongjie Wang, Theerawat Ruenkam, Rebecca E. Schelling, Vinayak Juvekar, Boby Mathew, Thorntun Deangphare, Wenxue Li, Yansheng Liu, Sarah A. Slavoff, Chayasith Uttamapinant, Ken H. Loh

## Abstract

Proximity labeling (PL) methods utilize spatially targeted chemical or enzymatic generation of a diffusible, reactive intermediate to covalently tag neighboring proteins in living systems. Unlike other tools for studying molecular interactions, PL can detect transient protein relationships with high spatial and temporal sensitivity, allowing for insight into their roles in biological processes. However, current enzymatic PL tools, such as TurboID and APEX2, are limited by their substrate structure and chemistry, which can generate significant background and/or perturb cellular physiology. To address these limitations, we have developed aminoacyl-tRNA synthetase ID (aaRSID), a PL tool that leverages an engineered pyrrolysyl tRNA synthetase (PylRS) for proximity labeling of proteins. We chose PylRS because it can catalyze promiscuous lysine labeling in the absence of its cognate tRNA and utilize a variety of non-canonical amino acids (ncAAs) as substrates. Here, we demonstrate aaRSID’s intrinsic proximity labeling activity, use directed evolution to improve this activity, and apply the improved mutant (aaRSID-Ma1.3) for subcellular proteomics and multiplexed imaging. Our work establishes aminoacyl-tRNA synthetases as a new PL enzyme class and introduces a versatile chemical platform for developing ncAA-derived probes to map cellular microenvironments, greatly expanding the applications possible of PL technology.

## Introduction

Proximity labeling (PL) has emerged as a powerful strategy for spatially mapping biomolecules and their interaction networks in living systems, with several enzyme platforms developed to generate reactive intermediates to covalently label biomolecules of interest within defined cellular environments. How intermediates are generated determines the practical trade-offs of each platform, including biological compatibility, spatial resolution and probe scope^1^. Peroxidase-based enzymes, including engineered ascorbate peroxidases (APEX/APEX2/APOX)^2–4^ and horseradish peroxidase (HRP)^5^, enable rapid labeling by oxidizing phenolic probes into short-lived radicals, but their reliance on exogenous H_2_O_2_ can impose oxidative stress and perturb cellular physiology. Recent O_2_-dependent platforms, such as tyrosinase (TyroID)^6^ and engineered laccase (LaccID)^7^, circumvent the need for exogenous H_2_O_2_, but are restricted to cell surface and extracellular proteome profiling. ATP-dependent PL enzymes based on small molecule-protein ligases, such as TurboID^8^, offer an orthogonal strategy by coupling ATP hydrolysis to generate reactive acyl-adenylate intermediates for protein labeling. Nevertheless, these biotin ligase-based platforms remain reliant on biotin as both the enzymatic substrate and the labeling handle, limiting flexibility in probe design. Biotin ligase-based platforms can also suffer from limited enrichment sensitivity arising from endogenous biotin and biotinylated proteins found in all cells, which can overwhelm the proximity labeling signal, especially when TurboID is targeted to a rare population of cells within a tissue (Fig. S1). Other ATP-dependent systems, such as PUP-IT^9^, move beyond biotin-based detection, but rely on the covalent transfer of a protein modifier rather than a compact small-molecule probe, imposing size restrictions. Together, these limitations highlight the need for new PL enzymes that operate through distinct chemical mechanisms and expand the accessible probe chemistry.

Aminoacyl-tRNA synthetases (aaRSs) represent another class of ATP-dependent enzymes whose reactive ester intermediates are energetically analogous to the biotinyl-adenylate generated by TurboID, and thus could potentially be repurposed for proximity labeling. During aaRS’s canonical reaction pathway, the catalytic domains of aaRSs use ATP to activate amino acids as reactive aminoacyl-adenylate intermediates, followed by transfer of the aminoacyl group to the 3’ terminus of cognate tRNAs. Importantly, advances in genetic code expansion (GCE) over the last 25 years have established archaeal^10^ pyrrolysyl-tRNA synthetase (PylRS) as a highly engineerable aaRS capable of accepting a broad range of noncanonical amino acid substrates (ncAAs). These include those ncAAs bearing bioorthogonal chemical handles that are generally water soluble and bioavailable, allowing them to penetrate tissue when administered in the drinking water of small rodents^11–13^. These ncAAs provide access to diverse chemistries for downstream conjugation and bioorthogonal enrichment, allowing for the choice of detection or enrichment tags to be tailored to the application. PylRS is highly orthogonal to prokaryotic and eukaryotic aaRSs and their cognate amino acid substrates, enabling its application in a wide range of cells^14, 15^. We therefore reasoned that redirecting ncAA-derived aminoacyl-adenylate intermediates generated by PylRS could provide a biologically distinct and chemically versatile strategy for protein proximity labeling.

Here, we leverage existing PylRS engineering to repurpose this enzyme for proximity labeling in a platform termed aminoacyl-tRNA synthetase ID (aaRSID). We found that PylRSs from the *Methanomethylophilus alvus* (*Ma*)^16, 17^ and *Methanogenic archaeon* ISO4-G1 (*G1*)^16, 18^ strains can catalyze promiscuous lysine aminoacylation in the absence of their cognate tRNA^Pyl^. This lysine-labeling activity of PylRS is mechanistically distinct from ncAA incorporation in GCE or BONCAT^19^: rather than conjugating ncAAs to cognate tRNAs for co-translational incorporation into nascent polypeptide chains through ribosomal translation, the activated ncAA-ester intermediate is released and modifies lysine sidechains of proximal proteins in one step, just like biotinylation with TurboID. This novel proximity labeling activity of PylRS is maintained across *E. coli*, yeast, and mammalian cells, and can utilize multiple ncAA substrates, many of which are commercially available. We further show that aaRSID labeling activity can be enhanced through directed evolution to generate an improved enzyme for proximity proteomics. By coupling the engineerability and broad substrate scope of PylRS with proximity labeling, aaRSID expands the probe chemistry accessible to enzyme-mediated PL.

## Results

### Characterization of PylRS proximity labeling activity

PylRS aminoacylates its cognate tRNA^Pyl^ through a two-step ATP-dependent mechanism, in which L-pyrrolysine is first activated as an aminoacyl-AMP intermediate and then transferred to tRNA^Pyl^ to form Pyl-tRNA^Pyl^. In addition to L-pyrrolysine, PylRS variants can accept diverse ncAAs and generate the corresponding aminoacyl-AMP analogs. We observed that in the absence of cognate tRNA^Pyl^, two previously reported archaeal PylRS enzymes, *Ma*PylRS and *G1*PylRS, instead promoted conjugation of the aminoacyl group to proteins in solution (Fig. 1a). *In vitro* assays using N-propargyl-L-lysine (PrK) as the probe showed that *Ma*PylRS and *G1*PylRS covalently labeled both themselves and bovine serum albumin (BSA) in a PrK- and ATP-dependent manner (Fig. 1b, c). The resulting alkyne-tagged products were detected by copper(I)-catalyzed azide-alkyne cycloaddition (CuAAC) with commercially available azide reporters, including biotin-azide, Cy5-azide, and fluorescein (FAM)-azide (Fig. 1b). We also tested a truncated *Methanosarcina mazei* (*Mm*) PylRS, commonly used in genetic code expansion studies^20, 21^. Based on prior studies, we removed the N-terminal domain (*Mm*PylRS^ΔN2–184^), which includes an in-frame nuclear localization sequence^22^. In contrast to *Ma*PylRS and *G1*PylRS, *Mm*PylRS^ΔN2–184^ showed minimal protein labeling in solution (Fig. 1d).

**Figure 1.**
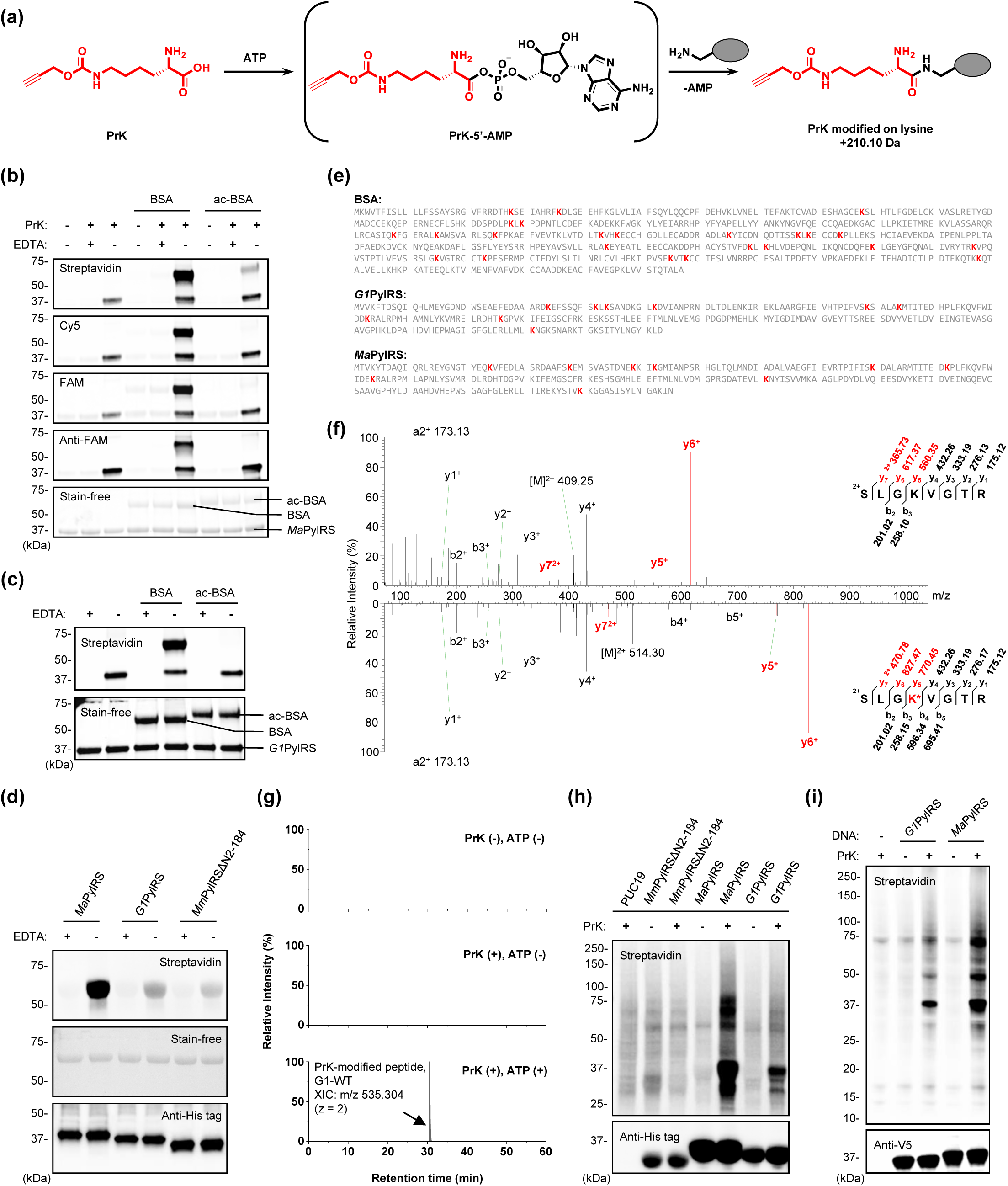
PylRS can catalyze tRNA-independent labeling of lysines. (a) Scheme describing PylRS-dependent adenylation of the ncAA PrK, which can tag primary amines on lysine resides. (b-c) *In vitro* assays showing purified *Ma*PylRS in (b) and *G1*PylRS in (c), can label BSA protein with PrK. This Mg^2+^-dependent reaction is inhibited by the addition of EDTA, or when lysine residues on BSA have been pre-acetylated (ac-BSA). PrK labeling is detected by click functionalization with biotin-azide followed by streptavidin detection (1^st^ row in b & upper panel in c), Cy5-azide or FAM-azide followed by fluorescence detection (2^nd^ and 3^rd^ rows in b, respectively), or FAM-azide followed by anti-FAM detection (4^th^ row in b). Total protein levels are detected in BioRad stain-free gels (5^th^ row in b & lower panel in c). (d) Comparison of PrK-dependent labeling by purified *Ma*PylRS, *G1*PylRS, and *Mm*PylRSΔN2-184 *in vitro*. Labeling was detected by click functionalization with biotin-azide followed by streptavidin detection, with EDTA-containing reactions included as negative controls. Total protein levels were detected by stain-free imaging, and PylRS enzymes were detected by anti-His-tag immunoblotting. (e) *In vitro*, PylRS-labeled BSA as in (b & c) was digested into peptides and analyzed by LC-MS/MS. Labeled peptides were detected on 24 sites on BSA, 9 sites on *G1*PylRS, and 9 sites on *Ma*PylRS. (f) Representative MS/MS spectra of an unlabeled BSA peptide (top row) and its PrK-labeled counterpart (bottom row), showing the mass shift associated with propargyl lysylation. (g) Extracted ion chromatograms (XICs) of the peptide (ac)SLGKVGTR modified with PrK (m/z 535.304) by *G1*PylRS. (h) Western blot showing promiscuous labeling in bacteria. PylRS variants (or PUC19 in the negative control) were overexpressed in *E. Coli* BL21(DE3) strain and PrK (or no PrK in the negative controls) was provided for 4 hours to initiate the labeling. PrK on labeled proteins was detected using click chemistry functionalization with biotin-azide, followed by streptavidin detection. (i) Western blot showing promiscuous labeling in mammalian cells. PylRS variants (or DNA minus in the negative control) were overexpressed in HEK293T cells and PrK (or no PrK in the negative controls) was provided overnight to initiate the labeling. PrK on labeled proteins was detected using click chemistry functionalization with biotin-azide, followed by streptavidin detection.

To validate lysine as the labeling site on proteins, we chemically blocked lysine ε-amines on BSA with excess N-succinimidyl acetate (acetyl-NHS) to generate acetylated BSA (ac-BSA). Lysine acetylation markedly reduced PrK-dependent labeling signals, indicating that free lysine ε-amines are required for PrK labeling (Fig. 1b, c). In a separate experiment without acetyl-NHS pretreatment, we identified 24 PrK-modified sites on BSA and 9 self-labeling sites on both *Ma*PylRS and *G1*PylRS by LC-MS/MS analysis following tryptic digestion of the labeled BSA to peptides (Fig. 1e). We then compared the MS/MS spectrum of a representative PrK-labeled peptide with that of its unmodified, backbone-matched counterpart (Fig. 1f). The corresponding mass shifts in the y5, y6, and y7 ions localized the modification to the lysine residue. Consistent with our previous assays, *in vitro* assays using a lysine-containing peptide as the substrate demonstrated PrK- and ATP-dependent labeling of the peptide (Fig. 1g). These experiments demonstrate that native PylRS enzymes exhibit proximity labeling activity with an ncAA *in vitro*, and that the mechanism of their proximity labeling chemistry – that is, release of the activated aminoacyl ester intermediate for reaction with nucleophilic lysine side chains in solution – is analogous to that of proximity biotinylation by TurboID as expected.

We next asked whether the proximity labeling activity we observed *in vitro* could be extended to the interior of living cells. To this end, we expressed *Ma*PylRS, *G1*PylRS, and *Mm*PylRS^ΔN2–184^ in both *E. coli* (Fig. 1h) and cultured human cells (Fig. 1i). We observed PrK-dependent proximity labeling by both *Ma*PylRS and *G1*PylRS in *E. coli* and HEK293T cells, with *Ma*PylRS consistently producing stronger labeling signals. Similar to the *in vitro* assay, we detected no proximity labeling with *Mm*PylRS^ΔN2–184^ in *E. coli*.

Together, these experiments demonstrate that homologs of PylRS have intrinsic activity for proximity labeling. In the absence of their cognate tRNA, *Ma*PylRS and *G1*PylRS can label lysine residues on proteins in solution in an ATP-dependent manner. Labeling occurs with PrK, an ncAA, suggesting that the labeling activity of PylRS is amenable to non-canonical substrates outside of L-pyrrolysine. Because the reactivity of aminoacyl adenylates is expected to be spatially restricted due to reaction with solvent, similar to biotin-adenylate, we hypothesized that certain PylRS homologs could act as a template for developing a new class of proximity labeling enzymes. We named this platform aminoacyl-tRNA synthetase ID, or aaRSID.

### Engineered PyIRS variants expand the substrate scope of aaRSID

Having established direct lysine modification with PrK, we next examined whether aaRSID could utilize additional ncAAs for proximity labeling. We selected five previously reported PylRS ncAA substrates bearing different bioorthogonal moieties, including the azide-containing amino acid N-((2-azidoethoxy)carbonyl)-L-lysine (NAEK), the strained alkene and alkyne amino acids norbornene-L-lysine (NBOK), trans-cyclooctene-L-lysine (TCOK), and exo-bicyclononyne-L-lysine (BCNK), and the photo-crosslinking amino acid propargyl-diazirine-L-lysine (PrDK)^23^ (Fig. 2a). These ncAAs provide diverse bioorthogonal “click” chemistry handles for CuAAC, tetrazine ligation (inverse-electron-demand Diels-Alder, IEDDA), strain promoted azide-alkyne cycloaddition (SPAAC), and photo-crosslinking, respectively^24^. For bulky ncAAs such as NBOK, TCOK, and BCNK, we tested proximity labeling using PylRS variants carrying mutations known to expand the substrate binding pocket to promote activation of these substrates. Among these, substitution of Y306 in *Mm*PylRS (corresponding to Y126 in *Ma*PylRS and Y125 in G1PylRS; sequence alignment in Fig. S2) with alanine, in combination with other mutations, has been widely used to generate polyspecific aaRSs for genetic code expansion^25–28^. LC-MS/MS analysis confirmed ATP-dependent formation of NAEK-, TCOK-, NBOK-, and BCNK-modified (ac)SLGKVGTR peptides by aaRSID (Fig. 2b). Substrate preferences differed markedly between PylRS variants. aaRSID-G1 efficiently transferred the smaller azide-containing substrate NAEK, whereas the bulkier TCOK, NBOK, and BCNK were detected only with the Y125A/M128L mutant (aaRSID-G1-AL)^28^. This is consistent with previous reports that substrate-binding pocket enlargement enables accommodation of bulkier ncAAs. This also confirms that the aaRSID approach can be generalized to labeling proteins with multiple bioorthogonal modalities by utilizing PylRS mutants with broadened ncAA substrate scopes, opening the possibilities for multiplexed and user-application tailored ncAA probes.

**Figure 2.**
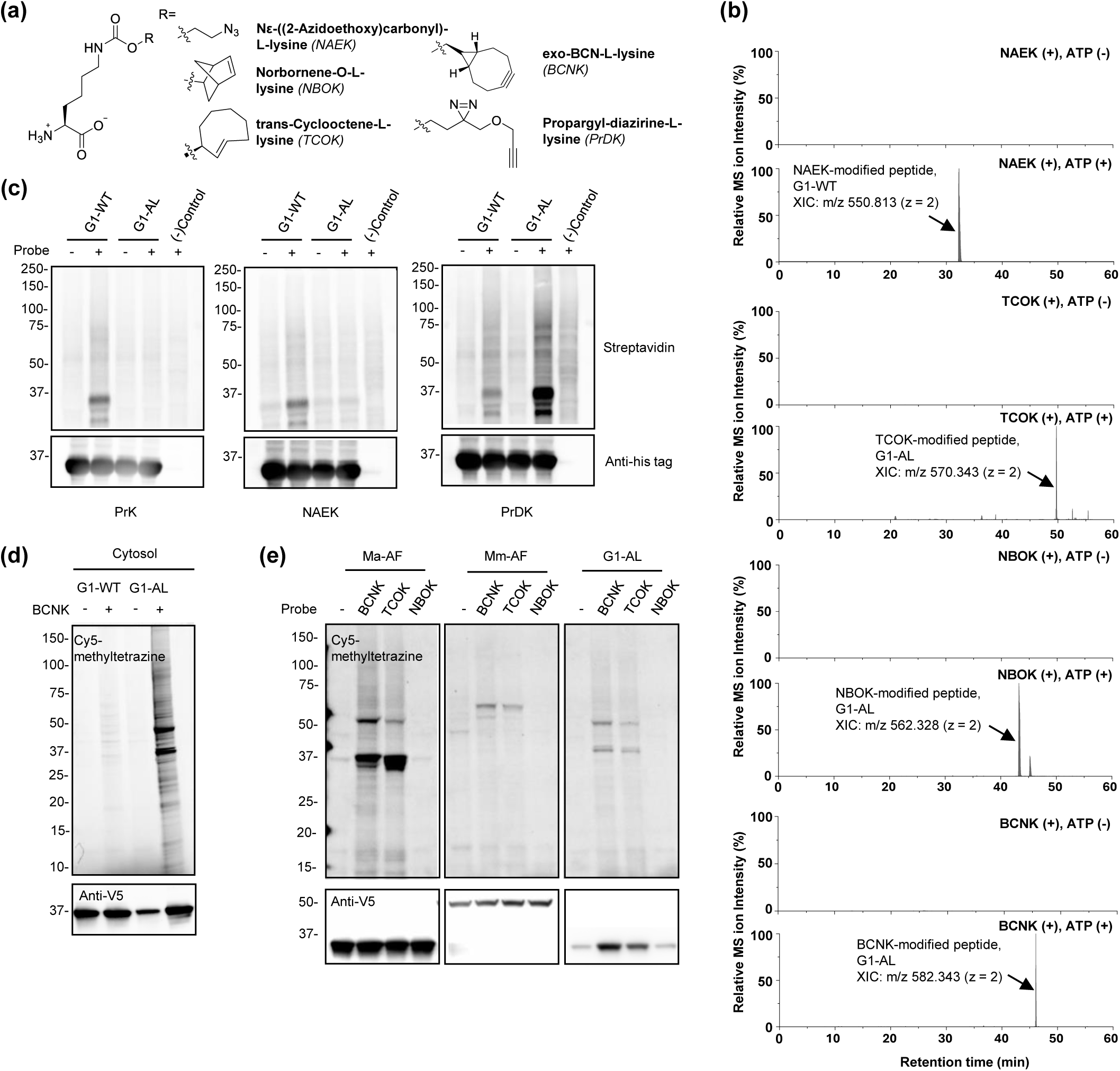
Specific PylRS mutants can catalyze proximity labeling with a wide variety of ncAA substrates. (a) ncAAs tested in this study. (b) XIC showing LC-MS/MS analysis of the peptide (ac)SLGKVGTR modified with NAEK (m/z 550.813) by G1PylRS (wt). The corresponding TCOK (m/z 570.343), NBOK (m/z 562.328) or BCNK (m/z 582.343) modified peptide was detected only when used with the (Y125A/M128L, “AL”) PylRS mutant. (c) Western blot showing proximity labeling with various ncAAs in bacteria using G1PylRS mutants. Similar to 1(g), PylRS (wt and Y125A/M128L mutants) were overexpressed in *E. Coli.* and ncAAs were provided overnight to initiate proximity labeling. PrK on labeled proteins was detected using click chemistry functionalization with biotin-azide, followed by streptavidin-HRP detection. NAEK was detected using click chemistry functionalization with biotin-alkyne. PrDK was detected using biotin-azide. (d) Western blot showing proximity labeling in mammalian cells by PylRS is dependent on which PylRS mutant was used. Similar to 1(h), PylRS (wt and Y125A/M128L mutants) were overexpressed in HEK293T cells and ncAAs were provided overnight to initiate proximity labeling. BCNK on labeled proteins was detected using click chemistry functionalization with Cy5-methyltetrazine, while anti-V5 detected expression levels of the enzyme. (e) Western blot showing proximity labeling in mammalian cells by PylRS with ncAAs depicted in 2(a) and detected similarly to (d).

We next examined the substrate preferences of aaRSID variants with expanded scope for proximity labeling in cells. In *E. coli*, aaRSID-G1 supported protein labeling with the relatively compact PrK and NAEK, whereas stronger labeling with the longer photo-crosslinking amino acid PrDK was detected predominantly in cells expressing aaRSID-G1-AL (Fig. 2c). The expanded substrate scope of aaRSID-G1-AL was also maintained in mammalian cells. HEK293T cells expressing aaRSID-G1 or aaRSID-G1-AL were incubated with BCNK, followed by Cy5-methyltetrazine derivatization. Consistent with the preference of aaRSID-G1-AL for bulkier ncAA substrates, BCNK labeling was observed only in aaRSID-G1-AL-expressing cells (Fig. 2d). We further compared several engineered PylRS scaffolds using strained alkene (TCOK) and alkyne (BCNK) substrates. Labeling efficiencies with these substrates differed among the enzymes, with aaRSID-Ma-AF (Y126A/Y206F)^27^ and aaRSID-G1-AL producing strong labeling, whereas aaRSID-Mm-AF (Y306A/Y384F)^25^ exhibited moderate activity (Fig. 2e).

These results demonstrate that prior substrate engineering of PylRS can be leveraged to generate distinct aaRSID scaffolds with orthogonal substrate specificities toward a wide variety of functionalized ncAAs, including multiple classes of “clickable” and photoreactive ncAAs. Furthermore, substrate specificity can be tuned through PyIRS engineering, expanding the range of chemical functionalities that can be introduced by aaRSID for downstream labeling and proteomic applications. The broad substrate compatibility of aaRSID suggested that further engineering could improve catalytic activity while preserving probe versatility.

### Directed evolution of aaRSID using yeast display

Although our preliminary results showed that aaRSID variants enabled proximity labeling with ncAAs in several contexts, their slow labeling required prolonged incubation (typically up to 24 h in mammalian cells) to generate detectable signals. We therefore explored both rational design and *in vitro* evolution in turn to improve the labeling efficiency of aaRSID. Inspired by the BirA R118G mutation, which transformed a highly specific biotin ligase into a promiscuous biotinylating enzyme^29^, we performed glycine-scanning mutagenesis of two conformationally dynamic regions in aaRSID-G1: the motif 2 loop and the β5–β6 hairpin. The motif 2 loop was selected based on structural studies of *Mm*PylRS showing that the corresponding region undergoes an open-to-closed conformational transition upon binding of AMP-PNP, a nonhydrolyzable ATP analog^30^. The β5–β6 hairpin, on the other hand, dynamically adopts open and closed conformations, repositioning Y204 between the exterior and interior of the amino acid-binding pocket^28^ and thereby suggesting a dynamic lid-like role over the active site. In parallel, we evaluated previously reported *Ma*PylRS variants evolved for PrK incorporation in GCE^31, 32^ for their ability to support proximity labeling. However, neither strategy yielded variants (45 mutants in total) with substantially enhanced labeling activity relative to the corresponding wild-type enzymes under the conditions tested (Fig. S3).

We therefore turned to yeast surface display coupled with fluorescence-activated cell sorting (FACS) for directed evolution^33^. This platform has been successfully used to engineer the catalytic efficiency of various PL enzymes, including biotin ligases (TurboID^8^ and ultraID^34^), peroxidases (APEX2^3^ and APOX^4^), and laccase (LaccID^7^). Furthermore, PylRS enzymes have previously been functionally expressed in *Saccharomyces cerevisiae* and engineered for genetic code expansion^32^, supporting yeast display as a suitable platform for evolving proximity PylRS variants. Given the lack of prior reports of surface-displayed PylRS mediating proximity labeling in yeast, we tested several wild-type aaRSID homologs (aaRSID-Ma, aaRSID-G1, and aaRSID-Mm) for their performance in the presence of PrK on the yeast cell surface. Surface-displayed aaRSID-Ma and aaRSID-G1 exhibited appreciable labeling activity under long PrK incubation times without prior engineering, thus readily serving as viable templates for directed evolution (Fig. S4a). Due to higher labeling activity of aaRSID-Ma, we selected this homolog for subsequent engineering (Fig. S4b).

We generated randomly mutagenized libraries of aaRSID-Ma by error-prone PCR and constructed a pooled library containing at least 10 variants, ranging from 0–7 mutations per gene (Fig. S5), which were displayed on the yeast surface through fusion to the Aga1p–Aga2p mating proteins. Labeling reactions were initiated by incubating cells with PrK, ATP, and MgCl_2_, followed by CuAAC derivatization with biotin picolyl azide^35^, which requires lower Cu^2+^ levels, improving cell health and viability for the selections. Labeling activity and surface expression were quantified using streptavidin-conjugated fluorophores and anti-Myc antibody staining, respectively (Fig. 3a). We reasoned that active aaRSID variants would generate a reactive aminoacyl-adenylate intermediate capable of labeling nearby surface lysines through a diffusion-or contact-dependent mechanism, or both. Selection stringency was progressively increased over successive rounds by shortening the labeling time from 6 h to 10 min and reducing the PrK concentration from 1 mM to 50 µM. After 11 rounds of selection spanning 3 generations, the population exhibiting high labeling-to-expression ratios was enriched by FACS (Fig. 3b, Fig. S6-10). Sequencing of the enriched clones identified mutations at five positions. The first-generation variant, aaRSID-Ma1.1, contained T20A, L104S, and F105L. The second-generation variant, aaRSID-Ma1.2, acquired K151E and N166H in addition to the three mutations of aaRSID-Ma1.1, whereas the final variant, aaRSID-Ma1.3, retained N166H but replaced K151E with K151G (Fig. 3c).

**Figure 3.**
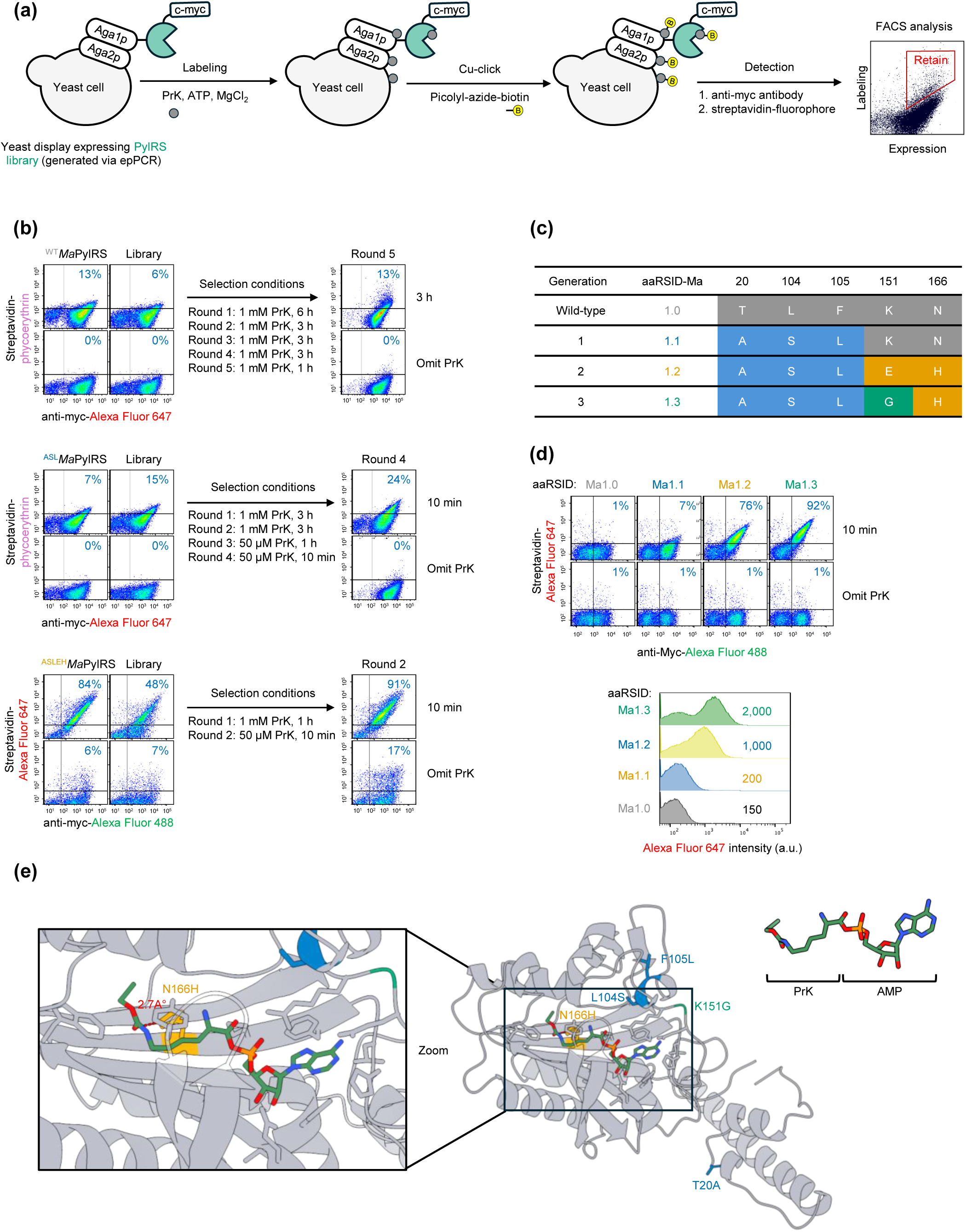
Yeast display-based directed evolution of aaRSID. (a) Schematic of the yeast surface display workflow for the directed evolution of aaRSID. Labeling activity and enzyme expression were detected by staining with streptavidin-phycoerythrin and anti-myc antibody, respectively. FACS was used to analyze and enrich cells exhibiting high labeling-to-expression ratios within the red trapezoid sorting gate. (b) Directed evolution campaign spanning 11 rounds of selection across three generations of evolution. Labeling conditions for each round are indicated below the arrows. Labeling activity was quantified as the percentage of PrK-labeling-positive cells (Q_2_) among total PylRS-expressing cells (Q_2_ + Q_4_). Experiments were performed independently twice with similar results. (c) Highly enriched mutations found in each generation. Wild-type amino acids are highlighted in grey, while other generations are highlighted in blue (generation 1), yellow (generation 2) and green (generation 3). (d) Comparison of individual aaRSID clones. Representative FACS plots and average Cy5 fluorescence intensity histograms of individual aaRSID clones following a 10-min labeling reaction. (e) Predicted structure of aaRSID-Ma1.3 bound to the PrK-AMP intermediate generated using Boltz-2 and visualized in ChimeraX. Representative accumulated residues are highlighted according to the color scheme shown in (c).

We next compared the evolved variants under the same labeling conditions. Relative to aaRSID-Ma1.0 (wild-type *Ma*PylRS), aaRSID-Ma1.1 exhibited ∼1.3-fold higher Alexa Fluor 647 labeling intensity. Labeling activity increased substantially following acquisition of N166H, with aaRSID-Ma1.2 and aaRSID-Ma1.3 exhibiting ∼6.7-fold and ∼13.3-fold higher Alexa Fluor 647 intensities, respectively (Fig. 3d, Fig. S11). Structural prediction of aaRSID-Ma1.3 in complex with the PrK-AMP intermediate using Boltz-2^36^ positioned H166 within the substrate-binding pocket, with its imidazole side chain in close proximity (∼2.7 Å) to the carbonyl oxygen of PrK-AMP, suggesting a potential hydrogen bond that may influence substrate recognition or positioning; the remaining evolved residues are positioned outside the substrate-binding pocket (Fig. 3e). These observations identify N166H as a critical mutation associated with the enhanced labeling activity observed and suggest that this improvement may be mediated by improved PrK binding.

### Characterization of the evolved aaRSID construct

Having identified evolved enzymes from the yeast display campaigns, we next validated if their enhanced labeling activity could be functionally transferred to other biological contexts. We first evaluated the variants in *E. coli* cells; we observed a labeling trend similar to that observed in our yeast evolution studies, with labeling activity (normalized to enzyme expression) increasing across successive generations of evolved variants. As anticipated, labeling activity in *E. coli* increased substantially in aaRSID-Ma1.2 and aaRSID-Ma1.3 (Fig. 4a), further supporting that N166H is the critical mutation for proximity labeling activity toward PrK. These results demonstrate that the enhanced labeling activity is maintained beyond the yeast display platform.

**Figure 4.**
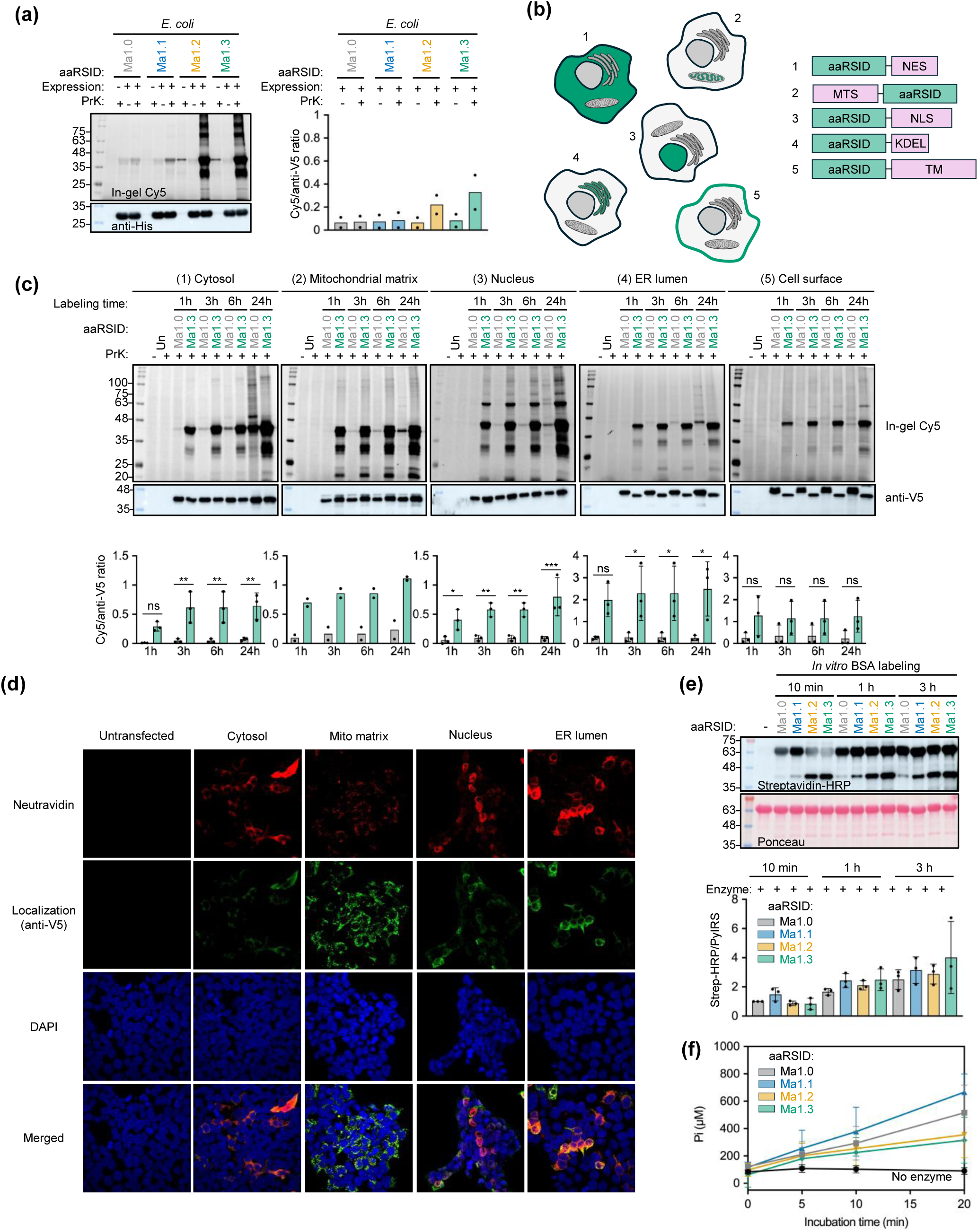
Characterization of evolved aaRSID enzymes. (a) Labeling activity of aaRSID variants in *E. coli*. Transformed cells were induced and labeled with PrK overnight. Whole-cell lysates were derivatized with Cy5-azide via CuAAC and analyzed by in-gel Cy5 fluorescence, while PylRS expression was detected by anti-His blotting. Cy5 fluorescence intensity was normalized to the corresponding anti-His signal (n = 2 independent experiments). (b) Subcellular targeting in HEK293T cells. aaRSID variants were directed to the cytosol (NES), ER lumen (KDEL), mitochondrial matrix (MTS), nucleus (NLS), or cell surface (TM) using the indicated targeting sequences. Targeted compartments are highlighted in green. (c) Time-course comparison of labeling activity between aaRSID-Ma1.0 and aaRSID-Ma1.3 across subcellular compartments in HEK293T cells. Transfected cells were labeled with PrK for 1, 3, 6, or 24 h. Whole-cell lysates were derivatized with Cy5-azid via CuAAC and analyzed by in-gel Cy5 fluorescence, while PylRS expression was detected by anti-V5 blotting. Cy5 fluorescence intensity was normalized to the corresponding anti-V5 signal (n = 2–3 independent experiments; error bars represent SD). Statistical significance was assessed by two-way ANOVA followed by Šídák’s multiple-comparisons test (\**P* < 0.05, \*\**P* < 0.01, \*\*\**P* < 0.001). Un, untransfected control. (d) Confocal fluorescence microscopy of HEK293T cells expressing compartment-targeted aaRSID-Ma1.3. Protein labeling was visualized by NeutrAvidin staining, PylRS expression and localization by anti-V5 staining, and nuclei by DAPI. (e) In vitro BSA labeling with PrK by purified aaRSID-Ma variants. BSA was incubated with the indicated variants in the presence of PrK and ATP for 10 min, 1 h, or 3 h. PrK-modified proteins were derivatized with biotin-azide via CuAAC and detected by streptavidin-HRP blotting, while total protein was visualized by Ponceau S staining. Relative labeling activity was quantified as the streptavidin-HRP signal normalized to the corresponding PylRS signal from Ponceau S staining (n = 2–3 independent experiments; error bars represent SD). -, no enzyme control. (f) Comparison of PrK-dependent adenylation activity of aaRSID variants. PrK-dependent pyrophosphate (PPi) production was monitored over time using a malachite green assay in the presence of excess PrK and a fixed ATP concentration (n = 3 independent experiments; error bars represent SD). PPi released during adenylation was converted to inorganic phosphate for colorimetric detection. A reaction lacking enzyme served as a negative control.

We then assessed the evolved enzymes in mammalian cells, a context most often used for proximity labeling-mediated spatial chemoproteomics. aaRSID variants were fused to subcellular localization signals or domains targeting the cytosol (NES), mitochondrial matrix (Mito), nucleus (NLS), endoplasmic reticulum (ER) lumen (KDEL), or the cell surface (TM) in HEK293T cells (Fig. 4b) as previously performed^6, 8^. Across all compartments, aaRSID-Ma1.2 and aaRSID-Ma1.3 consistently outperformed the wild-type enzyme and earlier-generation variants (Fig. 4c-f, Fig. S12), confirming that the enhanced labeling activity is broadly transferable across diverse intracellular and cell-surface environments. Because aaRSID-Ma1.3 consistently exhibited the highest labeling activity in both *E. coli* and HEK293T cells, we selected this variant for further characterization. Time-course analyses revealed substantially higher labeling activity than the wild-type enzyme in all intracellular compartments examined, with labeling detectable after 1 h of incubation and generally increasing over time. In contrast, cell-surface labeling showed no significant increase across the labeling time points, potentially owing to limited ATP stability under the extracellular labeling conditions and the normal absence of ATP in the extracellular space^37, 38^. We also noticed that aaRSID-Ma1.3 expression varied among subcellular compartments. Compared with the wild-type enzyme, aaRSID-Ma1.3 was expressed at lower levels in the cytosol, ER lumen, and on the cell surface, but accumulated to higher levels in the mitochondrial matrix and nucleus (Fig. 4c). Despite these differences in expression, aaRSID-Ma1.3 consistently exhibited enhanced labeling activity, indicating that its improved performance is not solely attributable to increased enzyme abundance.

To verify correct subcellular localization and labeling activity, we performed confocal immunofluorescence imaging of aaRSID-Ma1.3 targeted to each compartment. Following overnight incubation with PrK, robust neutravidin staining colocalized with anti-V5 immunofluorescence in all targeted compartments, confirming both the expected localization and labeling activity of aaRSID-Ma1.3 in living cells (Fig. 4d). aaRSID-Ma1.3 significantly outperformed the wild-type enzyme across all compartments tested (Fig. S13). Comparing these localized constructs to aaRSID-Ma1.3 without a subcellular localization signal also confirmed that the construct is not naturally sequestered to the nuclear compartment (Fig. S14), as has previously been reported for *Mm*PylRS containing its N-terminal domain^22^.

To gain insight into the labeling mechanism, we characterized the *in vitro* labeling activities of purified aaRSIDs. Labeling of BSA was detectable after 10 min of incubation and increased over time for all variants. However, in contrast to the trend observed in cell-based assays, the highly active aaRSID-Ma1.2 and aaRSID-Ma1.3 variants exhibited reduced BSA labeling at the earliest time point (10 min) relative to aaRSID-Ma1.1 and the wild-type enzyme. Notably, this reduced early intermolecular labeling coincided with progressively increased self-labeling of the evolved variants (Fig. 4e), suggesting altered partitioning of the reactive aminoacyl-adenylate intermediate between self-labeling and intermolecular substrate labeling. We then quantified the initial inorganic pyrophosphate (PPi) release rate, an indirect measure of the adenylation reaction, using a malachite green assay^39, 40^. Consistent with the early-time-point BSA-labeling results, aaRSID-Ma1.2 and aaRSID-Ma1.3 exhibited lower initial PPi release rates than aaRSID-Ma1.1 and aaRSID-Ma (Fig. 4f, Fig. S15). These findings indicate that the enhanced labeling activities of the later-generation variants observed in cells are not fully explained solely by increased adenylation activity. Instead, these results suggest that the cellular labeling results reflect catalytic steps beyond formation of the reactive intermediate, including its subsequent transfer to nearby acceptors, which are not captured by the simplified *in vitro* assays. Additional factors present in the cellular environment, such as protein proximity, molecular crowding, and transient protein interactions, may contribute to the superior labeling performance of the evolved variants, potentially by increasing effective local concentrations of acceptors and promoting more localized transfer of the reactive intermediate^41^.

### Application of aaRSID for proteomics

We next developed two distinct workflows for the enrichment and profiling of the PrK-labeled proteome in HEK293T cells expressing cytosolically targeted aaRSID-Ma1.3 (aaRSID1.3-NES). PrK-labeled proteins were conjugated by CuAAC to either biotin-azide for streptavidin enrichment, or FAM-azide for anti-FAM enrichment (Fig. 5a). Both workflows recovered aaRSID-dependent labeled proteins in the eluates following enrichment (Fig. 5b). Proteomic datasets were then filtered by receiver operating characteristic (ROC) analysis using annotated positive (cytosolic) and negative (nuclear) reference protein sets (Fig. S16a, b). The filtered datasets contained 1,048 and 851 cytosol-annotated proteins, respectively, with 680 proteins shared between workflows, representing 64.9% and 79.9% of the corresponding sets (Fig. 5d). This substantial overlap supports concordant recovery of cytosolic proteins through two distinct affinity-enrichment strategies.

**Figure 5.**
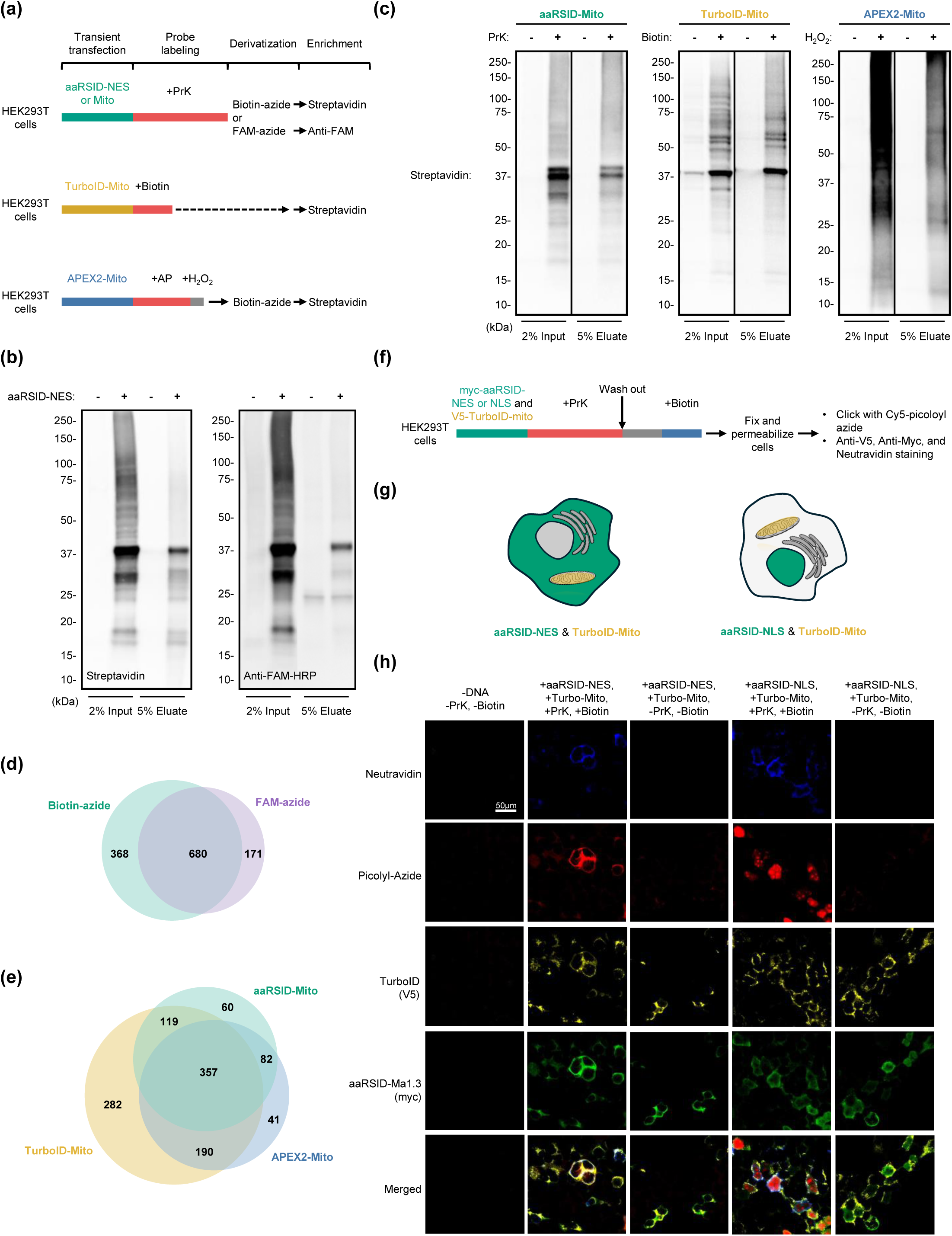
Application of aaRSID for proteomics and multiplexed protein labeling. (a) Scheme showing aaRSID-Ma1.3-catalyzed N-Propargyl-L-Lysine (PrK) labeling, TurboID-catalyzed biotinylation, and APEX2-catalyzed alkyne-phenol (AP) tagging in HEK293T cells, along with the corresponding enrichment workflows. (b, c) Representative blots of input and enriched fractions from the indicated proximity labeling systems. aaRSID-Ma1.3-NES samples were analyzed by streptavidin or anti-FAM-HRP blotting (b), whereas aaRSID-Ma1.3-Mito, TurboID-Mito and APEX2-Mito samples were analyzed by streptavidin blotting (c). (d) Venn diagram showing the overlap between cytosolic protein sets obtained using cytoplasm-targeted aaRSID (aaRSID-NES) with biotin-azide/streptavidin and FAM-azide/anti-FAM enrichment workflows. (e) Venn diagram showing the overlap among mitochondrial protein sets obtained using mitochondrial matrix-targeted aaRSID, TurboID, and APEX2. The biotin-azide/streptavidin workflow was used for aaRSID-Mito. (f∼h) Multiplexed labeling in mammalian cells using aaRSID and TurboID. (f) Labeling protocol used. (g) Schematic showing aaRSID targeted to the cytoplasm (aaRSID-NES) or nucleus (aaRSID-NLS) and TurboID targeted to the mitochondrial matrix (TurboID-Mito). (h) Confocal imaging of aaRSID- and TurboID-labeled proteins. Proteins labeled by aaRSID using PrK were detected by click conjugation with Cy5-picolyl-azide, while proteins biotinylated by TurboID were detected using NeutrAvidin.

We then extended the aaRSID-based proteomic profiling to mitochondria and compared the proteins identified using aaRSID with those identified using mitochondria-targeted TurboID (TurboID-Mito)^42^ and APEX2 (APEX2-Mito)^3^. PrK- and alkyne-phenol (AP)^43^-labeled proteins generated by aaRSID-Mito and APEX2-Mito, respectively, were conjugated to biotin-azide via CuAAC before streptavidin enrichment, whereas TurboID-biotinylated proteins were enriched directly with streptavidin (Fig. 5a). Following ROC-based filtering using annotated positive (mitochondrial matrix) and negative (cytosolic) reference protein sets (Fig. S16 c-e), the aaRSID-Mito dataset contained 618 proteins with mitochondrial annotations (Fig. 5e). Of these, 558 (90.3%) were also obtained using TurboID-Mito or APEX2-Mito, including 357 proteins shared by all three methods. This substantial overlap indicates that aaRSID-mediated PrK labeling enables the identification of many of the same mitochondrial proteins detected by established proximity-labeling methods.

### Orthogonal dual-modality proximity labeling with aaRSID and TurboID

While aaRSID and TurboID utilize similar chemistries for protein labeling, they use different substrates for downstream enrichment. We therefore reasoned that we could use the two labeling methods in tandem for multiplexing subcellular proteomic analysis in mammalian cells. To test this theory, we transiently expressed TurboID-Mito with either nuclear (NLS) or cytosolic (NES) aaRSID-Ma1.3 in HEK293T cells (Fig. 5g). Briefly, aaRSID labeling proceeded for 24 h followed by a 1 h washout step to remove excess PrK (Fig. 5f). 30 min prior to fixation, we initiated TurboID labeling with the addition of biotin. Cells were immediately fixed after labeling for immunofluorescence imaging. Localized, compartment-specific labeling with PrK was detected for both aaRSID-Ma1.3-NLS and aaRSID-Ma1.3-NES (Fig. 5h), demonstrating that aaRSID remained generally active regardless of the local cellular environment.

Our multiplexed labeling strategy resulted in robust orthogonal labeling in both TurboID/Ma1.3 pairs (Fig. 5h). As anticipated, labeling from both methods was confined to the subcellular localization of each enzyme, confirming the spatial specificity of aaRSID-Ma1.3 based on enzyme expression. Non-transfected cells, along with PrK- and biotin-negative controls, showed no labeling signal. Together, these data suggest that aaRSID-Ma1.3 is suitable for live cell labeling in conjunction with TurboID, enabling PL experiments across multiple compartments in a single cell.

### The expanded substrate scope for AL mutants is maintained in aaRSID-Ma1.3

Having established that the Y126A/M129L (AL) mutations expand the substrate scope of wild-type *Ma*PylRS toward bulky ncAAs, we next asked whether this broadened substrate specificity could be transferred to the evolved aaRSID-Ma1.3 scaffold. Because aaRSID-Ma1.3 was evolved using PrK as the selection substrate, it remained unclear whether the evolved scaffold would remain amenable to substrate-pocket engineering for structurally distinct ncAAs. We therefore introduced the AL mutations into aaRSID-Ma1.3 to generate aaRSID-Ma1.3-AL and examined whether the resulting variant retained the ability to utilize the bulky ncAAs TCOK and BCNK.

To assess the effects of the AL mutations on substrate preference, we compared aaRSID-Ma1.3 and aaRSID-Ma1.3-AL using the same *in vitro* peptide-labeling assay (as performed in Fig. 2b) with the standard peptide (ac)SLGKVGTR and four ncAA substrates: PrK, NAEK, TCOK, and BCNK. Extracted ion chromatograms showed ATP-dependent formation of PrK- and NAEK-modified peptides with aaRSID-Ma1.3, whereas little labeling was detected with TCOK or BCNK (Fig. 6a). Introducing the AL mutations shifted this substrate preference: aaRSID-Ma1.3-AL supported ATP-dependent peptide labeling with TCOK and BCNK, while signals corresponding to PrK- and NAEK-modified peptides were markedly reduced (Fig. 6b).

**Figure 6.**
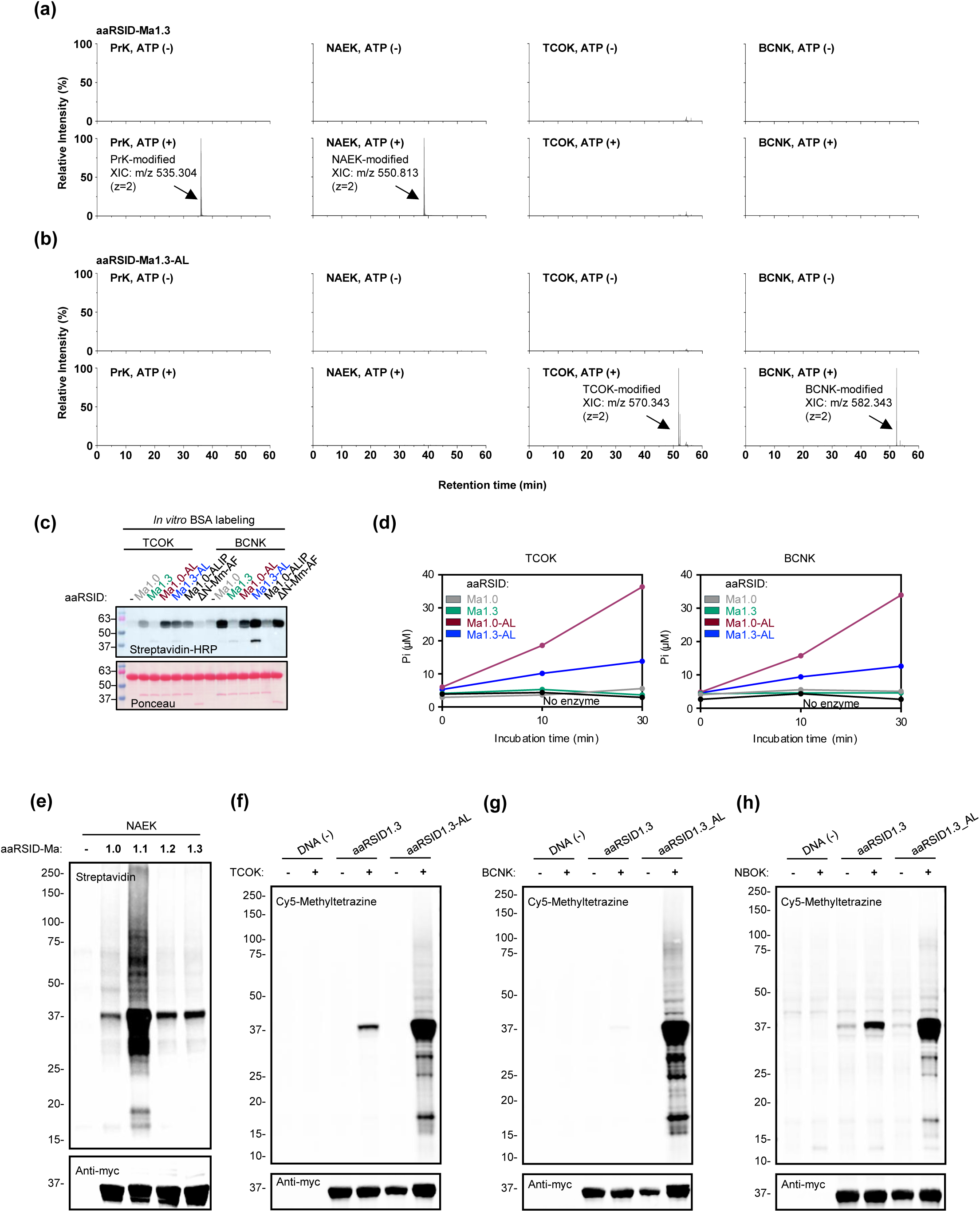
Expanded substrate scope of aaRSID-1.3 with the Y125A/M128L binding pocket mutations. (a, b) Extracted ion chromatograms (XICs) of the peptide (ac)SLGKVGTR modified with PrK (m/z 535.304), NAEK (m/z 550.813), TCOK (m/z 570.343), or BCNK (m/z 582.343) by aaRSID1.3 (a) or its Y125A/M128L mutant, aaRSID1.3-AL (b). Reactions performed with or without ATP were analyzed by LC–MS/MS. All indicated ions are doubly charged. For each substrate, intensities were normalized to the highest signal across the four XICs shown in (a) and (b), which was set to 100%. (c) In vitro BSA labeling of bulky ncAAs (TCOK and BCNK) by enlarged pocket PylRS variants. Labeling was analyzed after 1 h-incubation by streptavidin-HRP blotting; PylRS loading was assessed by Ponceau S staining. (d) Initial PPi release kinetics of bulky ncAAs (TCOK (left) and BCNK (right)) by aaRSIDs determined via malachite green assay. A reaction omitting the enzyme served as a negative control. (e) Streptavidin blotting showing NAEK labeling in cells expressing the indicated aaRSID-Ma1.X-NES variants. Following labeling, proteins were conjugated to biotin-alkyne via CuAAC. (f-h) In-blot fluorescence analysis of TCOK (f), BCNK (g), and NBOK (h) labeling in mammalian cells expressing aaRSID1.3-NES or aaRSID1.3-AL-NES. Following labeling, proteins were detected with Cy5-methyltetrazine.

We then tested whether this compatibility with bulky substrates extended to protein labeling using an *in vitro* BSA-labeling assay (Fig. 6c). For comparison, we included *Ma*PylRS-ALIP (Y126A/M129L/H227I/Y228P), previously engineered for efficient genetic incorporation of TCOK^27^, and *Mm*PylRS-AF^ΔN2–184^ (Y306A/Y384F, hereafter ΔN-*Mm*PylRS-AF), derived from a variant engineered for bulky ncAA incorporation^25^. With TCOK, aaRSID-Ma1.0-AL (Ma1.0 corresponding to wild-type *Ma*PylRS, here with added AL mutations) showed the strongest labeling activity, followed by aaRSID-Ma1.3-AL and *Ma*PylRS-ALIP, as determined by BSA labeling intensity normalized to aaRSID protein levels. In contrast, ΔN-*Mm*PylRS-AF showed little detectable labeling under these conditions. With BCNK, both aaRSID-Ma1.0-AL and aaRSID-Ma1.3-AL labeled BSA; *Ma*PylRS-ALIP showed little detectable activity while ΔN-*Mm*PylRS-AF supported detectable labeling. Unexpectedly, wild type aaRSID exhibited detectable BSA labeling with both bulky ncAAs despite lacking the AL mutations. Overall, the activity of aaRSID-Ma1.3-AL toward both TCOK and BCNK demonstrates that the expanded substrate compatibility conferred by the AL mutations is retained in the evolved aaRSID-Ma1.3 background.

We next examined whether the observed labeling activities were reflected at the substrate-activation step by monitoring PPi release using a malachite green assay (Fig. 6d). aaRSID-Ma1.0-AL exhibited the highest adenylate formation activity toward the bulky ncAAs (TCOK and BCNK), with PPi production increasing progressively over time. Importantly, aaRSID-Ma1.3-AL retained detectable PPi production with both substrates, whereas aaRSID-Ma1.3 lacking the AL mutations remained near background levels. These results demonstrate that introduction of the AL mutations enables the evolved aaRSID-Ma1.3 scaffold to adenylate bulky ncAA substrates for PL, although PPi production does not directly measure protein-proximity labeling efficiency. However, aaRSID-Ma1.3-AL exhibited lower activity than the aaRSID-Ma1.0-AL predecessor, indicating that the effects of the AL and aaRSID-Ma1.3 mutations are not fully independent. Wild-type aaRSID also remained near background in the adenylation assay (Fig. 6d), contrasting with its apparent BSA-labeling signal. Thus, the BSA-labeling and substrate-activation results for wild-type aaRSID were not concordant under the conditions tested, and the basis of its apparent labeling remains unresolved.

Finally, we examined ncAA-dependent protein proximity labeling in mammalian cells. We first compared NAEK labeling across the aaRSID-Ma1.0–Ma1.3-NES variants, with Ma1.0 corresponding to wild-type, in HEK293T cells, followed by CuAAC conjugation to biotin-alkyne and streptavidin blotting. aaRSID-Ma1.1 showed the strongest overall labeling signal, whereas labeling returned to near-WT levels in aaRSID-Ma1.2 and aaRSID-Ma1.3 (Fig. 6e). Thus, successive generations of evolution using PrK did not uniformly improve labeling with an alternative substrate. The reduction in NAEK labeling following the second generation of evolution suggests that mutations introduced at this stage, particularly N166H, may contribute to altered substrate specificity.

We also compared TCOK,BCNK, and NBOK labeling by aaRSID-Ma1.3-NES and aaRSID-Ma1.3-AL-NES in HEK293T cells. Following conjugation to Cy5-methyltetrazine, in-blot fluorescence analysis showed that aaRSID-Ma1.3-AL produced stronger labeling signals than aaRSID-Ma1.3 with all three substrates (Fig. 6f-h), consistent with the shift in substrate preference observed in the peptide-labeling assays (Fig. 6a-b).

Together, these results demonstrate that the substrate preference of PrK-evolved aaRSID-Ma1.3 can be redirected toward TCOK and BCNK through introduction of the AL mutations. Although the resulting aaRSID-Ma1.3-AL variant exhibited reduced adenylation activity (as measured by lower PPi production) with these bulky substrates relative to aaRSID-Ma1.0-AL, it retained the ability to activate and support protein labeling with bulky ncAAs that were poorly utilized by aaRSID-Ma1.3. Thus, directed evolution of aaRSID with PrK as the substrate may constrain activation of these bulky ncAAs but did not preclude subsequent engineering of its substrate scope to accommodate them. These findings suggest that the evolved aaRSID-Ma1.3 scaffold remains amenable to substrate-specific engineering and could provide a general framework for incorporating PylRS mutations recognizing different ncAAs. Overall, the distinct substrate preferences of aaRSID-Ma1.3 and aaRSID-Ma1.3-AL provide a starting point for developing orthogonal protein-labeling systems using different ncAA substrates.

## Discussion

aaRSID is a PylRS-based proximity labeling platform that redirects aminoacyl-tRNA synthetase chemistry toward tRNA-independent protein modification. A distinguishing feature of aaRSID is its chemical programmability inherited from PylRS, whose substrate recognition can be engineered to accommodate a diverse range of ncAAs normally used in genetic code expansion. We showed that aaRSID can install different bioorthogonal functionalities onto proteins, allowing the proximity labeling reaction to be decoupled from downstream detection or enrichment. The evolved aaRSID-Ma1.3 further enabled labeling across multiple cellular compartments and recovered subcellular compartment-enriched proteomes that substantially overlapped those obtained with established PL systems. Moreover, its substrate diversity allowed aaRSID to operate alongside TurboID for orthogonal labeling within the same cell. The utility of orthogonal PL chemistries has been demonstrated by approaches such as TransitID, which combine sequential labeling systems to resolve proteins trafficking between cellular compartments^44^. Thus, aaRSID provides an engineerable enzyme-substrate framework in which substrate recognition and transferred chemical functionality can be diversified, offering opportunities to develop additional orthogonal labeling channels for multiplexed and potential dynamic spatial proteomic studies. This includes pulse chase experiments often difficult to execute with biotin-ligase based PL because of the detrimental impacts on cellular physiology under biotin starvation.

aaRSID-Ma1.3 was generated from the intrinsic protein labeling activity of *Ma*PylRS through yeast display directed evolution, substantially improving labeling in cellular environments. Interestingly, this improvement did not correlate with increased amino acid adenylation or early *in vitro* BSA labeling, suggesting that cellular proximity labeling activity is not determined solely by aminoacyl-AMP formation. Productive labeling may instead depend on downstream processes, including transient enzyme-protein encounters or local transfer of the activated intermediate. This contact-dependent mechanism remains to be established but has been recently investigated for TurboID^41^. Importantly, introduction of the Y126A/M129L enlarging substrate-pocket mutations into aaRSID-Ma1.3 enables labeling with bulky ncAAs such as TCOK and BCNK, showing that the evolved scaffold remains amenable to substrate-specific engineering. Further evolution could therefore improve labeling efficiency while generating aaRSID variants with distinct ncAA preferences, providing a route toward chemically orthogonal PL systems.

## Limitations of the study

Despite demonstrated enhanced activity of aaRSID-Ma1.3 over the wild-type enzyme across different cell types and subcellular compartments, which we hypothesize is a result of promiscuous reaction of the aminoacyl adenylate intermediate with proximal proteins, the precise mechanism underlying its tRNA-independent proximity protein labeling remains to be investigated. Although aaRSID recovered compartment-enriched proteomes that substantially overlapped those obtained with established PL systems, its labeling radius and spatial resolution have not yet been directly determined. Only a small fraction of the ncAA repertoire accessible to engineered PylRS systems has been explored, which may significantly influence these factors in a context-dependent manner. Finally, the present study was limited to cultured cellular systems; the feasibility of aaRSID in animal models remains to be established, particularly with respect to ncAA delivery, labeling efficiency, and bioorthogonality in complex tissues. Further optimization of labeling kinetics and substrate specificity, together with *in vivo* validation, will therefore be important for extending aaRSID toward spatial and multiplexed proteomic applications in more complex biological systems.

## Acknowledgements

Funding: K.L., Z.W., R.S., V.J., B.M., were supported by funding from the NIH (P30DA018343 and R35GM162138), the American Heart Association, the Esther A. & Joseph Klingenstein Fund and the Simons Foundation and Oak Ridge Associated Universities. C.U. acknowledges funding support from VISTEC and the NSRF via the Research and Innovation Acceleration Agency for Competitiveness and Area Development (RCAD) (Program Management Unit for Technology and Innovation for Future Industries (PMU-B): Brainpower for Future Industries) [grant number B38G690002]. R.S. was also supported by NIH predoctoral training grants (T32GM149444, T32GM156537). T.R. and T.D. are supported by PhD studentship funds from VISTEC. S.S. was supported in part by a Mark Foundation for Cancer Research Emerging Leader Award, a Paul G. Allen Frontiers Group Distinguished Investigator Award, and a Sloan Research Fellowship (FG-2022-18417).

## Methods

### Overexpression and purification of evolved aaRSID mutants

aaRSID-Ma variants cloned in a pET21a bacterial expression vector were heat-shock transformed into competent BL21 (DE3) *E. coli* cells. Transformed *E. coli* cells were plated onto LB agar plates containing 100 µg/mL ampicillin and incubated overnight at 37 °C. Single colonies were picked to inoculate overnight starter cultures in ampicillin-supplemented LB medium at 37 °C with shaking. Saturated starter cultures were diluted 1:100 into 1 L of LB medium containing 100 µg/mL ampicillin and grown at 37 °C until reaching an OD_600_ of 0.6–0.8. Protein expression was induced ∼20 h at 16 °C by adding 0.7 mM isopropyl β-D-1-thiogalactopyranoside (IPTG). Cells were harvested by centrifugation at 15,000×g for 20 min at 4 °C and resuspended in ice-cold binding buffer (20 mM sodium phosphate, 300 mM NaCl, 10 mM imidazole, pH 7.4) supplemented with 1% Halt^TM^ protease inhibitor cocktail and 1 mM PMSF. Cells were disrupted via sonication on ice, and lysates were clarified by centrifugation at 15,000×g for 30 min at 4 °C. Gravity-flow columns containing Chelating Sepharose Fast Flow resin (Cytiva) were prepared manually and charged overnight with 0.2 M NiSO_4_ prior to use. Columns were equilibrated with binding buffer before loading the clarified protein lysates. The columns were washed sequentially with binding buffer and wash buffer (binding buffer supplemented with 25 mM imidazole). Target proteins were eluted using elution buffer (binding buffer supplemented with 300 mM imidazole). Purified proteins were concentrated and buffer-exchanged into storage buffer (100 mM HEPES, pH 7.5, 100 mM NaCl, 10 mM MgCl_2_, 4 mM DTT, and 20% glycerol) using a Vivaspin 20 centrifugal concentrator with a 10 kDa molecular weight cut-off (MWCO, Cytiva). Protein concentrations were determined by a BCA assay, and purity was verified via 12% SDS-PAGE analysis.

### *In vitro* labeling activity of purified variants

aaRSID variants (1 µg) were incubated in a 100 mM Na-HEPES buffer containing 1 mM PrK, 1 mM ATP, 10 mM MgCl_2_, and 30 µg of BSA at 37°C for designated intervals (10 min, 1 h, and 3 h). Reactions were terminated at each time point by snap-freezing in liquid nitrogen. For bulky ncAAs (TCOK and BCNK), labeling reactions were performed under the same conditions analyzed after 1 h. Following labeling, 10 µg of protein from each reaction was derivatized with biotin-azide via CuAAC (for PrK) or tetrazine-biotin (for TCOK and BCNK) via IEDDA, followed by streptavidin blotting as described below. PylRS loading was assessed by Ponceau S staining.

### Adenylation activity measurements

Adenylation activities were determined using a malachite green method following previously published protocols^39, 40^. Reaction mixtures were prepared in a 200 mM HEPES-K (pH 7.5) buffer containing 4 mM DTT, 10 mM MgCl_2_, 0.2 mM ATP, 10 mM ncAA, 4 U/mL *E. coli* inorganic pyrophosphatase (NEB), and 2.5 µM purified protein variants. Reactions were incubated at 37°C. At designated time points (PrK: 0, 5, 10, and 20 min; BCNK or TCOK: 0, 10, and 30 min), 10 µL aliquots were removed and immediately quenched on ice by resuspending in an equal volume of 20 mM EDTA (pH 8.0). Each aliquot was diluted with 140 µL of 200 mM HEPES-K, followed by the addition of 40 µL of an in-house prepared malachite green/molybdate reagent. The colorimetric mixtures were incubated at room temperature for 30 min. Absorbance at 620 nm was measured on an Infinite M Plex multimode microplate reader (Tecan). Raw absorbance values were converted to phosphate concentration using a KH_2_PO_4_ standard calibration curve (0-100 µM) (Fig. S15).

### Yeast cell culture

*Saccharomyces cerevisiae* strain EBY100 cells (ATCC) were cultured according to established protocols^8^. Cells were propagated at 30°C in yeast extract peptone dextrose (YPD) medium. The pCTCON2 yeast display plasmid encoding aaRSID-Ma variants, containing a *TRP* selection marker, was transformed into chemically competent yeast cells using the Frozen-EZ yeast transformation II kit (Zymo Research), following the manufacturer’s instructions. Transformants were then selected on synthetic dextrose plus casein amino acid (SDCAA) agar lacking tryptophan for 3 days. Single colonies were inoculated into 5 mL of SDCAA medium and grown overnight at 30°C with shaking. The resulting yeast culture was passaged twice prior to induction. To induce protein expression, saturated cultures were diluted at 1:9 ratio into SGCAA medium (SDCAA medium wherein dextrose is replaced with galactose) and incubated at 30°C with shaking for ∼20 h.

### Yeast display analysis and sorting by FACS

Induced yeast cells (2×10^6^ cells for individual clones; 1×10^7^ cells for libraries) were harvested and washed with PBS containing 0.1% BSA (PBS-B). Cells were then incubated in 3% BSA in PBS supplemented with 1 mM PrK, 1 mM ATP, and 10 mM MgCl_2_ for 10 min - 6 h at 30 °C with rotation. Following two washes with PBS, copper-catalyzed azide-alkyne cycloaddition (CuAAC) was performed at room temperature for 10 min using a reaction cocktail consisting of 10 μM CuSO_4_, 50 μM THPTA, 2.5 mM sodium ascorbate, 100 μM TEMPOL, and 10 μM biotin picolyl azide (Vector Labs) in PBS. After click labeling, cells were washed twice with PBS-B and incubated with a mouse anti-Myc primary antibody (1:500 dilution) for 30 min at room temperature with rotation. The cells were washed again with PBS-B and counter-stained on ice for 30 min with an anti-mouse secondary antibody conjugated to Alexa Fluor 647 (1:500 dilution, Invitrogen) or 488 (1:500 dilution, Abcam), and streptavidin-phycoerythrin (1:500 dilution, Jackson ImmunoResearch) or streptavidin-Alexa Fluor 647 (1:500 dilution, Invitrogen), respectively. Following a final wash, cells were resuspended in 500 μL of PBS-B.

Samples were analyzed or sorted on a BD FACSymphony S6 cell sorter (BD Biosciences) using the following configuration: a 561 nm excitation laser with a 586/15 emission filter for phycoerythrin; a 488 nm excitation laser with a 515/20 emission filter for AF488; and a 633 nm excitation laser with a 670/30 emission filter for Cy5 or Alexa Fluor 647. Individual clone comparisons were analyzed on a BD FACSMelody cell sorter (BD Biosciences) equipped with a 488 nm laser (527/35 emission filter for Alexa Fluor 488) and a 640 nm excitation laser (660/10 emission filter for Cy5 or Alexa Fluor 647). Data were recorded for 50,000-100,000 events per sample and analyzed using FlowJo^TM^ v10 software. Target populations were isolated using sequential gating strategies around the clustered population as follows: initial cells were plotted on a forward-scatter area (FSC-A) versus side-scatter area (SSC-A) to yield population 1 (P1). P1 was then resolved on a side-scatter width (SSC-W) versus side-scatter height (SSC-H) plot to isolate population 2 (P2). P2 was plotted on a forward-scatter width (FSC-W) versus forward-scatter height (FSC-H) plot to yield the single-cell population 3 (P3). P3 single cell population were then plotted using the X-axis for protein expression (anti-Myc) and the Y-axis for labeling activity (streptavidin conjugate).

For library selection, cells exhibiting the highest activity-to-expression ratios were gated and then sorted directly into a 50 mL conical tube containing 5 mL of SDCAA medium supplemented with 1% penicillin-streptomycin, 100 μg/mL carbenicillin and 50 μg/mL kanamycin. Collected cultures were subsequently incubated at 30 °C with shaking for 1–2 days. To identify mutations, a 1 mL aliquot of the expanded culture was miniprepped using the Zymoprep Yeast Plasmid Miniprep II kit (Zymo Research) according to the manufacturer’s instructions. The pooled plasmids were then transformed into competent DH5α *E. coli* cells (NEB) to isolate individual clones. At least 20 individual colonies were randomly picked and sequenced by Nanopore whole-plasmid sequencing (Quintara Biosciences). The remaining yeast culture was passaged using an inoculum of at least 10-fold excess of the total number of collected cells to initiate the subsequent round of selection.

### Yeast display directed evolution of aaRSID

aaRSID-Ma libraries were generated via error-prone PCR following a previously published protocol^8^. For each generation, 100 ng of template in pCTCON2 yeast display vector was amplified for 10-20 cycles using 0.4 µM forward and reverse primers flanking the aaRSID-Ma gene, 2 mM MgCl_2_, 5 units of Taq polymerase (NEB), and 2-20 µM each of the mutagenic nucleotide analogs 8-oxo-2′-deoxyguanosine-5′-triphosphate (8-oxo-dGTP, Jena Bioscience), and 2′-deoxy-P-nucleoside-5′-triphosphate (dPTP, Jena Bioscience). Mutagenized PCR products from each condition were gel-purified and further re-amplified in a scale-up volume of 200 µL standard PCR reaction to increase the yield, followed by a final gel purification step. The mutagenized insert (4 µg) was mixed with the *Bam*HI-*Nhe*I digested pCTCON2 vector (1 µg) and concentrated to a final volume of 10 µL using a speed vacuum concentrator. This DNA mixture was then electroporated into 400 µL of freshly prepared electrocompetent EBY100 yeast cells using a MicroPulser Electroporator (Bio-Rad). The electroporated cells were immediately recovered in 4 mL of YPD medium at 30°C for 30 min without shaking, followed by an additional 30 min with shaking. A 10 µL aliquot was serially diluted 100×, 1,000×, 10,000× and 100,000×, and 40 µL of each dilution was plated onto SDCAA agar at 30°C for 3 days. The remaining recovered culture was outgrowth in 100 mL SDCAA supplemented with 1% penicillin-streptomycin, 100 µg/mL carbenicillin and 50 µg/mL kanamycin at 30°C with shaking for 2 days. After 3 days, library sizes were calculated from the resulting colonies on the dilution plates, corresponding to 10^4^, 10^5^, 10^6^ or 10^7^ transformants, respectively.

### Generation 1

Three distinct libraries were generated using the wild-type *Ma*PylRS (aaRSID-Ma) as a template for error-prone PCR. Mutation rates were varied using the following conditions:

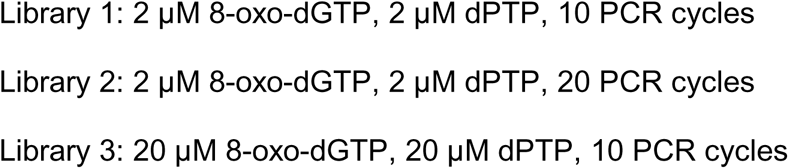

The resulting library sizes (calculated by the number of transformants as described above) were 2.2×10^7^, 2.1×10^7^ and 1.1×10^7^, respectively. FACS analysis showed robust protein expression in libraries 1 and 2, whereas library 3 exhibited poor expression, likely due to excessively high mutation rates. Therefore, libraries 1 and 2 were pooled to form the initial population for the first selection round. This combined library size had a total size of 4.3×10^7^ variants with an average of 1.3 amino acid changes per gene (range: 0-5 mutations).

Stringency was increased across 5 rounds of selection by reducing the labeling time from 6 h to 1 h. Labeled cells were derivatized with biotin picolyl azide via CuAAC and stained with an anti-Myc primary antibody, followed by an Alexa Fluor 647 secondary conjugate and streptavidin-phycoerythrin as detailed above. Selection parameters for each round are summarized below:

|  | Cells prepared | Labeling conditions | Cells sorted | Cells collected |
| --- | --- | --- | --- | --- |
| Round 1 | $5 \times 10^8$ | 1 mM PrK, 1 mM ATP, 10 mM MgCl<sub>2</sub>, 6 h | $1.8 \times 10^7$ | $1 \times 10^6$ (0.8%) |
| Round 2 | $2 \times 10^8$ | 1 mM PrK, 1 mM ATP, 10 mM MgCl<sub>2</sub>, 3 h | $1 \times 10^7$ | $5 \times 10^4$ (0.5%) |
| Round 3 | $1 \times 10^8$ | 1 mM PrK, 1 mM ATP, 10 mM MgCl<sub>2</sub>, 3 h | $8 \times 10^6$ | $5 \times 10^4$ (0.6%) |
| Round 4 | $1 \times 10^7$ | 1 mM PrK, 1 mM ATP, 10 mM MgCl<sub>2</sub>, 3 h | $5 \times 10^5$ | $2.5 \times 10^3$ (0.5%) |
| Round 5 | $5 \times 10^6$ | 1 mM PrK, 1 mM ATP, 10 mM MgCl<sub>2</sub>, 1 h | $1 \times 10^6$ | $2 \times 10^3$ (0.2%) |

Following each round, at least 20 individual colonies were randomly sequenced. Unique variants converging on consensus residues were screened on the yeast surface to compare their labeling activity against the wild-type enzyme. The double-mutant L104S/F105L and triple-mutant T20A/L104S/F105L (aaRSID-Ma1.1) displayed comparable labeling activities that significantly surpassed the wild-type enzyme. We selected both mutants as templates for the next generation of directed evolution.

### Generation 2

To optimize mutation frequencies, the concentration of mutagenic nucleotide analogs was adjusted to 4 µM, as the 20 µM concentration utilized in Generation 1 severely impaired protein expression. Six libraries were constructed via error-prone PCR using either the SL or ASL mutants as templates under the following conditions:

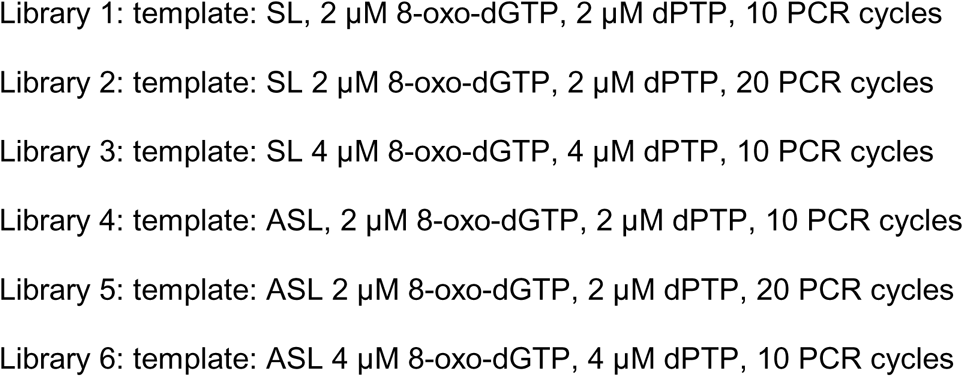

The library sizes were 7.5×10^5^, 2.3×10^5^, 2.6×10^6^, 1.5×10^6^, 1.8×10^6^ and 3.7×10^6^, respectively. FACS analysis confirmed robust expression across all six libraries. The libraries were combined as an initial population of ∼1×10^7^ variants with an average of 2.2 amino acid changes per gene (range: 0-7 mutations).

In this generation, selection stringency was increased by shortening incubation times (from 3 h to 10 min) and reducing the substrate concentration from 1 mM to 50 µM PrK across 4 rounds. CuAAC derivatization and antibody staining protocols remained identical to Generation 1. Selection round configurations are outlined below:

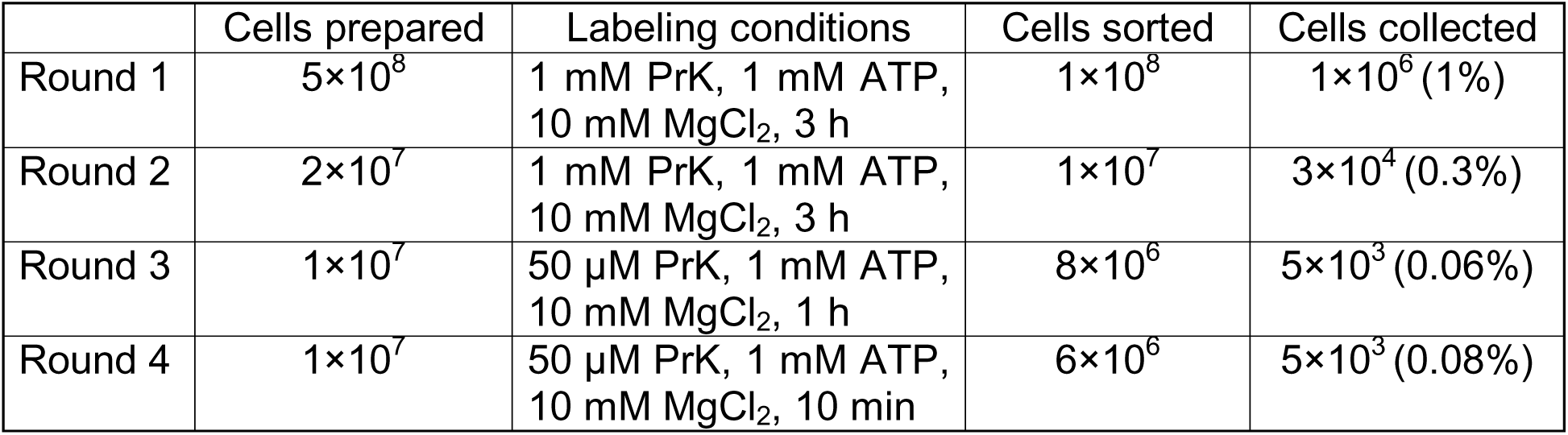

|  | Cells prepared | Labeling conditions | Cells sorted | Cells collected |
| --- | --- | --- | --- | --- |
| Round 1 | $5 \times 10^8$ | 1 mM PrK, 1 mM ATP, 10 mM MgCl<sub>2</sub>, 3 h | $1 \times 10^8$ | $1 \times 10^6$ (1%) |
| Round 2 | $2 \times 10^7$ | 1 mM PrK, 1 mM ATP, 10 mM MgCl<sub>2</sub>, 3 h | $1 \times 10^7$ | $3 \times 10^4$ (0.3%) |
| Round 3 | $1 \times 10^7$ | 50 μM PrK, 1 mM ATP, 10 mM MgCl<sub>2</sub>, 1 h | $8 \times 10^6$ | $5 \times 10^3$ (0.06%) |
| Round 4 | $1 \times 10^7$ | 50 μM PrK, 1 mM ATP, 10 mM MgCl<sub>2</sub>, 10 min | $6 \times 10^6$ | $5 \times 10^3$ (0.08%) |

Sequence analysis and clonal characterization on the yeast surface identified a prominent variant, T20A/L104S/F105L/K151E/N166H (aaRSID-Ma1.2), which markedly outperformed wild-type and aaRSID-Ma1.1 variants. Thus, the aaRSID-Ma1.2 variant was chosen as the parental template for Generation 3.

### Generation 3

Three libraries were generated using the aaRSID-Ma1.2 mutant as a template for error-prone PCR under the following conditions:

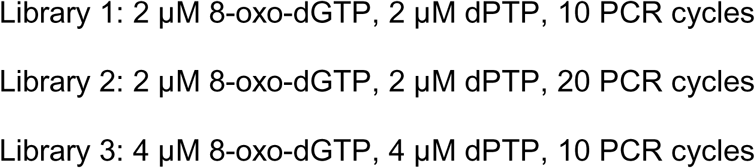

The library sizes were 8×10^6^, 4.5×10^6^ and 2.5×10^6^, respectively. Three libraries were combined as an initial population of 7.8×10^6^ with an average of 2 amino acid changes per gene (range: 0-7 mutations).

The selection strategy mirrored the stringent conditions used in Generation 2 with the details summarized in the table below:

|  | Cells prepared | Labeling conditions | Cells sorted | Cells collected |
| --- | --- | --- | --- | --- |
| Round 1 | $5 \times 10^8$ | 1 mM PrK, 1 mM ATP,<br>10 mM MgCl<sub>2</sub>, 3 h | $1 \times 10^8$ | $1 \times 10^6$ (1%) |
| Round 2 | $2 \times 10^7$ | 50 μM PrK, 1 mM ATP,<br>10 mM MgCl<sub>2</sub>, 3 h | $1 \times 10^7$ | $3 \times 10^4$ (0.3%) |

Sequencing of the Round 2 population revealed complete dominance by a single variant: T20A/L104S/F105L/K151G/N166H (aaRSID-Ma1.3).

After the Generation 3 selection had been completed, the PE detection channel of the flow cytometer became unreliable. This technical issue occurred after sorting and therefore did not affect the selection itself. For subsequent clone characterization and side-by-side comparison of variants across generations, PrK labeling was instead detected with streptavidin-Alexa Fluor 647, and PylRS surface expression was detected with an Alexa Fluor 488-conjugated secondary antibody.

Side-by-side surface display validation showed that the labeling activity of the aaRSID-Ma1.3 variant was equivalent to its parental aaRSID-Ma1.2 variant. Consequently, the directed evolution campaign was concluded, and both variants were transitioned to functional validation.

### Mammalian cell culture

HEK293T cells (ATCC) were cultured in complete DMEM media (gibco) supplemented with FBS (Corning) and penicillin-streptomycin (gibco) at 37 °C under 5% CO_2_. Cells were typically passaged at 90% confluency at a 1:10 dilution. For imaging in mammalian cell compartments, cells were seeded at ∼65% confluency on glass coverslips in 12-well plates. To improve the adherence of HEK293T cells, glass coverslips were pre-treated with 1mg/mL fibronectin (EMD Millipore) for >1 h at 37 °C before cell plating.

### Proximity labeling with aaRSID in mammalian cells

HEK293T cells (ATCC) were seeded onto a pre-coated 12 well plate and incubated at 37 °C overnight. Upon reaching ∼80% confluency, the cells were transfected with 900 ng pLJM1 plasmid encoding aaRSID variants fused to 5 different localization signals (NES, KDEL, MTS, NLS and TM) using PEI Max in Opti-MEM for 4h. The transfection medium was later exchanged to complete medium and further incubated overnight. Labeling was initiated by replacing the medium with complete medium supplemented with 1 mM PrK for designated time-course (1, 3, 6 or 24 h). Cells with TM-localized aaRSID also received a supplement of ATP and Mg^2+^. Prior to cell harvesting at each time point, the labeling medium was replaced with substrate-free complete medium for 1 h to permit the efflux of un-reacted free PrK from the intracellular space. Labeling activity was analyzed by in-gel Cy5 fluorescence or streptavidin western blotting, while enzyme expression was analyzed by anti-V5 western blotting, as described below.

### Proximity labeling with aaRSID in bacterial cells

aaRSID variants cloned into a pET21a bacterial expression vector were heat-shock transformed into competent BL21 (DE3) *E. coli* cells. Transformed *E. coli* cells were plated onto LB agar plates containing 100 µg/mL ampicillin and incubated overnight at 37 °C. Single colonies were randomly picked to inoculate overnight starter cultures in ampicillin-supplemented LB medium at 37 °C with shaking. Saturated starter cultures were diluted 1:100 into 1 L of LB medium containing 100 µg/mL ampicillin and grown at 37 °C until reaching an OD_600_ of 0.6–0.8. Protein expression was induced and labeled ∼20 h at 16 °C by adding 0.7 mM isopropyl β-D-1-thiogalactopyranoside (IPTG) and 1 mM PrK. Labeling activities and protein expression were analyzed by in-gel Cy5 fluorescence and western blotting, respectively, as described below.

### Gels and western blots

In brief, after live-cell labeling, cells were lysed on ice for 20 min with RIPA (for mammalian cells) or B-PER (for bacterial cells) buffer supplemented with 1% Halt^TM^ protease inhibitor cocktail and 1 mM PMSF. Lysates were then clarified by centrifugation at 18,000×g for 20 min at 4°C, and total protein concentrations were quantified using BCA assay (Sigma-Aldrich). 10 µg of protein lysate was incubated with a CuAAC click reaction mixture containing 200 µM CuSO_4_, 1 mM THPTA, 5 mM sodium ascorbate and 20 µM Cy5-azide or 20 µM biotin-azide for 1 h at room temperature. Reactions were quenched by adding SDS-loading dye followed by heating at 95°C for 5 min. Prepared samples were resolved on a 12% SDS-PAGE gel at 150 V for 70 min in a Tris-Glycine-SDS running buffer. To detect labeling activity, in-gel Cy5 fluorescence was visualized on Cy5 channel using an Amersham^TM^ ImageQuant^TM^ 800 imager (Cytiva). To determine protein expression, the gel was later transferred onto a 0.45 µm nitrocellulose membrane for 90 min at 350 mA. Total protein loading was assessed by staining the membrane with a Ponceau S solution followed by rinsing several times with DI water. Membranes were then blocked with 3% BSA in Tris-buffered saline containing 0.1% Tween-20 (TBST) for 1 h at room temperature and then incubated overnight at 4 °C with a mouse anti-V5 primary antibody (1:1,000 dilution, Cell Signaling) for mammalian constructs or a mouse anti-His primary antibody (1:1,000 dilution, Cell Signaling) for bacterial constructs in blocking buffer. Following three washes with TBST, membranes were incubated with an anti-mouse horseradish peroxidase (HRP)-conjugated secondary antibody (1:10,000 dilution, Bio-Rad) in blocking buffer for 1 h at room temperature, washed three times with TBST, and developed via chemiluminescence under an Amersham^TM^ ImageQuant^TM^ 800 imager (Cytiva).

### Immunofluorescence imaging

For aaRSID imaging in mammalian cell compartments, HEK293T cells were seeded on fibronectin-coated glass coverslips in 12-well plates. HEK293T cells expressing aaRSID-Ma or an aaRSID variant were prepared through overnight PEI transfection of aaRSID-encoding plasmids. Briefly, 2.15 µL PEI and 860 ng of plasmid were combined in 100 µL of serum-free media and added to cells for an overnight incubation. Cells were used for labeling experiments on the following day (∼18 h). To perform labeling, cell culture media was replaced with complete media containing 250 µM propargyllysine (PrK, SiChem) and incubated at 37 °C for 24 h or 2 h. For negative controls, we omitted PrK or the aaRSID enzyme. To remove excess PrK, PrK-containing media was replaced with complete DMEM 1 h prior to labeling cessation. After incubation, cells were washed 2 times with 1 mL DPBS and incubated in ice-cold methanol at −20 °C for 15 min to induce fixation and permeabilization. For the aaRSID localization experiments, cells were fixed and permeabilized after overnight transfection.

For the multiplexed imaging experiments with TurboID, HEK293T cells were cultured and seeded on glass coverslips as described above. Cells were transiently transfected with 645 ng of aaRSID-Ma1.3-NES or aaRSID-Ma1.3-NLS and 645 ng of TurboID-Mito overnight (∼18 h). For aaRSID labeling, cell culture media was replaced with complete media containing 250 µM PrK the following day and incubated for 24 h. Excess PrK was removed 1 h prior to labeling cessation by replacing the PrK-containing media. TurboID labeling was initiated by adding 250 µM biotin (Sigma-Aldrich) to the cells for 30 min before fixation. To stop the biotinylation reaction, cells were washed in ice-cold DPBS 3 times and kept on ice for 5 min. Cells were then incubated in 4% paraformaldehyde (Electron Microscopy Sciences) for 15 min at room temperature for fixation followed by 15 min in 0.5% Triton-X for permeabilization (Sigma-Aldrich).

Following permeabilization in both experiments described above, cells were blocked in 3% BSA (Sigma-Aldrich) in PBS-T (0.1% Tween 20, Thermo Scientific) for 1 h at room temperature. aaRSID-Ma1.3-labeled proteins were then derivatized using copper-catalyzed click chemistry. Briefly, a solution of 200 µM CuSO_4_, 1 mM BTTP, 2.5 mM sodium ascorbate, and either 100 µM biotin-azide (Vector Laboratories, Ma1.3-only imaging) or Cy5-picolyl-azide (Vector Labs, multiplexed imaging) was added to the cells. Plates were covered in aluminum foil and placed on a room temperature rocker. After one hour, cells were washed 3 times with PBS-T and incubated in mouse anti-Myc (Cell Signaling, 1:2000; aaRSID) and goat anti-V5 (Bethyl, 1:2000; TurboID) primary antibody overnight at 4 °C. The following morning, cells were again washed 3 times in PBS-T and incubated in secondary antibody at room temperature for 1 h. For the aaRSID-only experiments, cells were treated with donkey anti-mouse Alexa Fluor 488 (Abcam, 1:2000) and Neutravidin DyLight 594 (Invitrogen, 1:2000). Cells from the multiplexing experiment were treated with donkey anti-goat Alexa Fluor 568 (Invitrogen), donkey anti-mouse Alexa Fluor 488, and Neutravidin Dylight 405 (Invitrogen, 1:2000). Finally, the glass coverslips were washed in PBS-T, dried, and mounted onto glass slides (Electron Microscopy Sciences) with DAPI-containing (aaRSID-Ma1.3 only, SouthernBiotech) or DAPI-free (multiplexing, Invitrogen) mounting solution.

### Cell labeling for proteomics

HEK293T cells were seeded in poly-D-lysine-coated six-well plates. At 70–80% confluency, cells were transfected with 2.5 μg pcDNA3.1 plasmid encoding aaRSID-NES, aaRSID-Mito, TurboID-Mito, or APEX2-Mito using 6.25 μL PEI transfection reagent per well overnight.

For aaRSID-mediated labeling, the transfection medium was replaced with complete DMEM containing 1 mM PrK, and cells were incubated for 24 h. For aaRSID-NES experiments, cells receiving no plasmid DNA but treated with PrK served as negative controls. For aaRSID-Mito experiments, cells expressing the same construct without PrK treatment served as negative controls. Labeling was terminated by placing the plates on ice and washing the cells five times with ice-cold DPBS.

For TurboID-mediated labeling, cells expressing TurboID-Mito were incubated with complete DMEM containing 500 μM biotin for 15 min. Control cells were processed without supplemental biotin. Labeling was terminated by placing the plates on ice and washing the cells five times with ice-cold DPBS.

For APEX2-mediated labeling, cells expressing APEX2-Mito were pre-incubated with 500 μM alkyne-phenol (AP) at 37 °C for 2 h. Labeling was initiated by adding H_2_O_2_ to a final concentration of 1 mM for 1 min. The reaction was terminated by replacing the labeling medium with quenching solution containing 10 mM sodium ascorbate, 10 mM sodium azide, and 5 mM Trolox in PBS, followed by 3 washes with the same solution. Control cells were treated with AP without H_2_O_2_.

### Sample preparation for proteomics

Cells prepared as described above were harvested by scraping in ice-cold DPBS and pelleted by centrifugation at 300 × g for 5 min at 4 °C. Cell pellets were lysed in RIPA buffer (50 mM Tris-HCl, pH 8.0, 150 mM NaCl, 0.1% SDS, 1% Triton X-100, and 0.5% sodium deoxycholate) supplemented with 1× protease inhibitor cocktail by sonication. Lysates were clarified by centrifugation at 18,000 × g for 15 min at 4 °C.

Proteins were precipitated by sequential addition of methanol, chloroform, and water at a lysate:methanol:chloroform:water volume ratio of 1:4:1:3, with vortexing after each addition. Samples were centrifuged at 18,000 × g for 5 min, and the upper aqueous phase was removed. Precipitated proteins were washed three times with methanol, with centrifugation at 18,000 × g for 5 min between washes. Pellets were briefly air-dried and resuspended in PBS containing 1% SDS with sonication. Protein concentrations were determined using the Pierce BCA Protein Assay Kit.

For aaRSID and APEX2 samples, aliquots containing 1 mg of total protein were subjected to click reaction with 100 μM biotin-azide or fluorescein-azide, 1 mM CuSO_4_, 6 mM BTTP, and 10 mM sodium ascorbate in amine-free RIPA buffer (1 × PBS, 0.1% SDS, 1% Triton X-100, and 0.5% sodium deoxycholate). Reactions were incubated for 1 h at room temperature. Proteins were subsequently precipitated as described above, resuspended in PBS containing 1% SDS, and quantified by BCA assay.

For all three labeling methods (aaRSID, APEX2, and TurboID), equal amounts of protein from the labeled and corresponding control samples were used for enrichment based on BCA measurements. Protein samples in PBS containing 1% SDS were diluted with 5 volumes of SDS-free 2 × RIPA buffer (2 × PBS, 2% Triton X-100, and 1% sodium deoxycholate) and 4 volumes of water to obtain a final buffer composition of 0.1% SDS, 1% Triton X-100, and 0.5% sodium deoxycholate.

### Streptavidin enrichment protocol

Streptavidin enrichment of biotin-derivatized samples was performed as previously described^45^ with minor modifications. Specifically, 250 μL of streptavidin-coated magnetic bead slurry (Invitrogen) was washed twice with 1 mL of RIPA buffer and incubated with the protein samples described above overnight at 4 °C with end-over-end rotation. The beads were sequentially washed twice with 1 mL of RIPA buffer, once with 1 mL of 1 M KCl, once with 1 mL of 0.1 M Na_2_CO_3_, once with 1 mL 2 M urea in 10 mM Tris-HCl (pH 8.0), and twice with 1 mL of RIPA buffer.

For Western blot analysis, 5% of the bead suspension was collected during the final wash. Bound proteins were eluted by heating at 95 °C for 15 min in 2 × SDS loading buffer containing 20 mM DTT and 2 mM biotin. For proteomic analysis, the remaining 95% of the beads were washed once with 1 mL of 50 mM Tris-HCl (pH 7.5) and twice with 1 mL of 2 M urea in 50 mM Tris-HCl (pH 7.5). The beads were resuspended in 80 μL of 2 M urea in 50 mM Tris-HCl (pH 7.5) containing 1 mM DTT, 0.5 mM CaCl_2_, and 0.8 μg of trypsin (Promega), and incubated at 25 °C for 1.5 h with vigorous shaking. Following this initial digestion, the supernatant was collected, and the beads were washed twice with 60 μL of 2 M urea in 50 mM Tris-HCl (pH 7.5). The washes were pooled with the initial supernatant.

The pooled eluates were reduced with 8 mM DTT (final concentration) at 65 °C for 20 min and alkylated with 20 mM iodoacetamide (final concentration) at 37 °C for 30 min in the dark. The samples were diluted with 50 mM Tris-HCl (pH 7.5) to a final urea concentration of 1 M, and an additional 0.5 μg of trypsin was added. Digestion was continued overnight at 25 °C with shaking. Peptides were desalted using Pierce C18 Spin Columns according to the manufacturer’s instructions.

### Anti-fluorescein enrichment protocol

Anti-fluorescein enrichment of fluorescein-derivatized samples was performed as previously described^44^ with minor modifications. Specifically, 250 μL of Dynabeads Protein G slurry (Invitrogen) was washed twice with 1 mL of RIPA buffer and incubated with 30 μL of anti-fluorescein antibody (Abcam) in 1 mL of RIPA buffer for 1 h at 30 °C with end-over-end rotation. Antibody-bound beads were washed twice with 1 mL of RIPA buffer and incubated with the protein samples described above overnight at 4 °C with end-over-end rotation.

Beads were washed twice with 1 mL of RIPA buffer, twice with 1 mL of high-salt buffer (50 mM Tris-HCl, pH 7.5, 1 M NaCl, 1 mM EDTA, 0.1% SDS, 1% NP-40, and 0.5% sodium deoxycholate), and twice more with 1 mL of RIPA buffer. Bound proteins were eluted twice with 50 μL of elution buffer (50 mM Tris-HCl, pH 8.0, 150 mM NaCl, 1% SDS, 1% Triton X-100, and 0.5% sodium deoxycholate) at 50 °C for 20 min with vigorous shaking. Eluates were combined, and 5 μL was reserved for Western blot analysis. The remaining 95 μL was processed for protein clean-up and tryptic digestion using S-Trap micro columns (ProtiFi) according to the manufacturer’s instructions.

### Liquid chromatography and mass spectrometry

LC-MS/MS analysis of the cytosolic proteome and PrK-labeled peptides derived from BSA, G1PylRS, and *Ma*PylRS was performed on a Vanquish Neo UHPLC system (Thermo Scientific) connected to an Orbitrap Eclipse Tribrid mass spectrometer (Thermo Scientific) equipped with a nano-electrospray ion source. Data acquisition was controlled by Xcalibur (Thermo Scientific). Peptides were separated on a self-packed CoAnn column (75 μm inner diameter × 20 cm length) packed with ReproSil-Pur 120 Å C18-Q resin (1.9 μm particle size; Dr. Maisch GmbH, Ammerbuch, Germany). Solvent A consisted of 0.1% formic acid in water, and solvent B consisted of 80% acetonitrile containing 0.1% formic acid. Peptides were eluted using a 120-min gradient: 6-27% B over 90 min, 27-45% B over 10 min, 45-100% B over 10 min, followed by a 10-min hold at 100% B. The flow rate was maintained at 300 nL/min. The mass spectrometer was operated in data-dependent acquisition (DDA) mode. MS1 scans were acquired over an m/z range of 350–1400 at a resolution of 120,000 at m/z 200, with a normalized AGC target of 250% and the maximum injection time set to Auto. Following each MS1 scan, up to 20 of the most intense eligible precursor ions were selected for HCD fragmentation using an isolation window of 1.4 m/z and a normalized collision energy of 30%. MS2 spectra were acquired at a resolution of 30,000 at m/z 200, with the AGC target set to Standard and the maximum injection time set to Auto. The MS2 scan range was determined automatically. Dynamic exclusion was set to 20 s. The MS1 and MS2 spectra were recorded in profile and centroid modes, respectively. For analysis of the standard peptide (ac)SLGKVGTR labeled with ncAAs, a 60-min LC gradient was used: 6-27% B over 45 min, 27-45% B over 5 min, 45-100% B over 5 min, followed by a 5-min hold at 100% B. Dynamic exclusion was disabled. All other parameters remained unchanged.

The mitochondrial samples were measured using the DIA mass spectrometry method described previously^46–48^. LC separation was performed on EASY-nLC 1200 systems (Thermo Scientific, San Jose, CA) using a self-packed analytical PicoFrit column (New Objective, Woburn, MA, USA) (75 µm × 50 cm length) using C18 material of ReproSil-Pur 120A C18-Q 1.9 µm (Dr. Maisch GmbH, Ammerbuch, Germany). A 120-min measurement with buffer B (80% acetonitrile containing 0.1% formic acid) from 5% to 37% and corresponding buffer A (0.1% formic acid) during the gradient was used to elute peptides from the LC. The flow rate was kept at 300 nl/min, temperature-controlled at 60 °C using a column oven (PRSO-V1, Sonation GmbH, Biberach, Germany). The Orbitrap Fusion Lumos Tribrid mass spectrometer (Thermo Scientific) instrument coupled to a nanoelectrospray ion source (NanoFlex, Thermo Scientific) was calibrated using Tune (version 3.0) instrument control software. Spray voltage was set to 2,000 V and heating capillary temperature at 275 °C. The DIA-MS methods consisted of one MS1 scan and 33 MS2 scans of variable isolated windows. The MS1 scan range is 350-1650 m/z, and the MS1 resolution is 120,000 at m/z 200. The MS1 full scan AGC target value was 2.0E6, and the maximum injection time was 50 ms. The MS2 resolution was set to 30,000 at m/z 200 with the MS2 scan range 200-1800 m/z, and the normalized HCD collision energy was 28%. The MS2 AGC was set to be 1.5E6, and the maximum injection time was 50 ms. The default peptide charge state was set to 2. Both MS1 and MS2 spectra were recorded in profile mode.

### Mass spectrometry data processing

For identification of PrK-labeled peptides derived from BSA, G1PylRS, and MaPylRS, DDA-MS data were searched using MaxQuant version 2.4.10.0 against a custom FASTA database containing only the protein sequences present in each reaction. Unless otherwise specified, default search parameters were used. Propargyl lysylation (+210.100442 Da) on lysine with a neutral loss of 56.026215 Da was included as an additional variable modification.

The DDA-MS data from cytosolic proteome experiments were searched against the Swiss-Prot human protein database (downloaded on February 24, 2026, containing 20,431 entries) using MaxQuant version 2.4.10.0. Unless otherwise specified, default search parameters were used, and label-free quantification (LFQ) was enabled. Mass tolerances were set to 4.5 ppm for MS1 precursor matching in the main search and 20 ppm for MS2 matching. Trypsin/P was selected as the protease with a maximum of 2 missed cleavages allowed. The digestion mode was set to Specific. Carbamidomethylation of cysteine was selected as a fixed modification. Oxidation of methionine and protein N-terminal acetylation were selected as variable modifications, allowing a maximum of 5 variable modifications per peptide. False discovery rates (FDRs) at the peptide-spectrum match (PSM) and protein levels were controlled at 1% using a target-decoy approach. The site decoy fraction was set to 1%. Match between runs was enabled. For cytosolic proteome analysis, protein groups flagged as “Only identified by site”, “Reverse”, or “Potential contaminant” were removed from the dataset.

The DIA-MS data were analyzed using DIA-NN version 2.3.2 in library-free mode^49^ against the Swiss-Prot human protein database (downloaded on February 24, 2026, containing 20,431 entries). Unless otherwise specified, default search parameters were used. Trypsin/P was selected as the protease with a maximum of 2 missed cleavages allowed. Carbamidomethylation of cysteine was specified as a fixed modification. N-term M excision was specified as a variable modification. Mass accuracy was automatically optimized, and match between runs was enabled. The precursor and protein FDR threshold was set to 1%.

The unfiltered lists of identified proteins were first processed using Perseus version 2.1.5.0. Protein intensities were log2-transformed. Proteins with valid values in all replicates of at least one experimental group (either the labeled group or the negative control group) were retained. Proteins were further required to have at least two unique peptides (MaxQuant) or two distinct proteotypic peptide sequences (DIA-NN). Missing values were imputed in Perseus using random values drawn from a normal distribution, with a width of 0.3 and a down shift of 1.8.

### ROC-based filtering of cytosolic proteomic data

The filtered proteins are shown in Table S1. The true-positive (TP) reference set comprised cytosolic proteins annotated with the Gene Ontology (GO) term GO:0005829. The false-positive (FP) reference set comprised nuclear proteins annotated with any of the following GO terms^8^: GO:0016604, GO:0031965, GO:0016607, GO:0005730, GO:0001650, GO:0005654, GO:0005634. Proteins included in the TP reference set were excluded from the FP reference set. For each protein, log_2_(fold change) was calculated by subtracting the mean log2 intensity of the negative-control replicates from that of the labeled replicates. Log_2_(fold change) values for all proteins were then normalized by subtracting the median log_2_(fold change) of the FP reference proteins, setting the FP median to 0. To determine the log_2_(fold change) threshold, receiver operating characteristic (ROC) analysis was performed. Proteins were ranked in descending order of normalized log_2_(fold change). For each protein on the ranked list, the cumulative TP and FP counts above its normalized log_2_(fold change) value were calculated. These counts were divided by the respective total numbers of TP and FP reference proteins in the filtered dataset to obtain the true-positive rate (TPR) and false-positive rate (FPR). An ROC curve was generated by plotting TPR against FPR. The cutoff was selected to maximize TPR-FPR (Fig. S16).

### ROC-based filtering of mitochondrial proteomic data

The filtered proteins are shown in Table S1. Data normalization and ROC-based filtering were performed as described above for the cytosolic proteomic data, using different TP and FP reference sets. The true-positive (TP) reference set comprised mitochondrial matrix proteins annotated with GO:0005759, excluding proteins annotated with any of the following GO terms: GO:0005741 (mitochondrial outer membrane), GO:0005758 (mitochondrial intermembrane space), and GO:0005743 (mitochondrial inner membrane)^44^. The false-positive (FP) reference set comprised cytosolic proteins annotated with GO:0005829. Proteins included in the TP reference set were excluded from the FP reference set.

### Proximity labeling with TurboID in mice

TurboID x CD4+ mice were generated and treated as previously described^50^. Briefly, Biotin-containing water was prepared by dissolving solid biotin in sterile drinking water at a concentration of 0.25 mg/mL. Mice were provided *ad libitum* access to biotin-containing water during treatment. Following treatment, mice were anesthetized with an overdose of isoflurane and perfused with PBS to minimize any non-specific residual biotinylation signal. Organs were then dissected, rinsed with ice-cold PBS briefly, snap frozen in liquid nitrogen, and stored at −80°C before use.

---

**Supplementary Figure 1.**
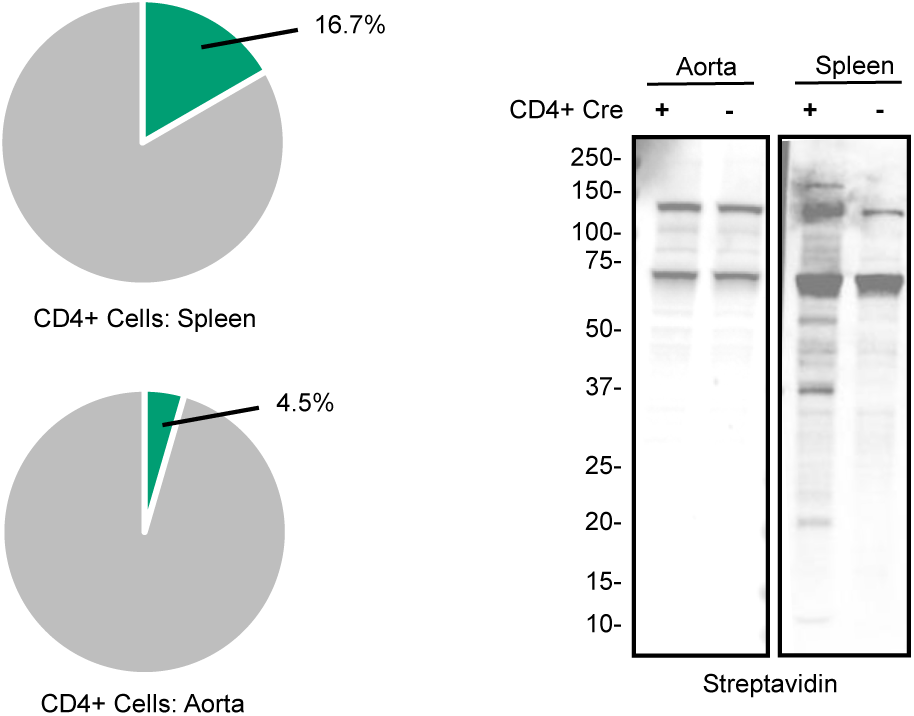
Background information supporting the development of aaRSID. CD4+ x TurboID mice were fed biotin water alongside cre-controls for two days to initiate proximity labeling. Tissue with a high (spleen) and low (aorta) percentage of CD4+ cells were collected for enrichment of TurboID-labeled proteins. Biotinylated proteins from homogenized tissue were visualized via western blot using IRDye 800CW Streptavidin.

**Supplementary Figure 2.**
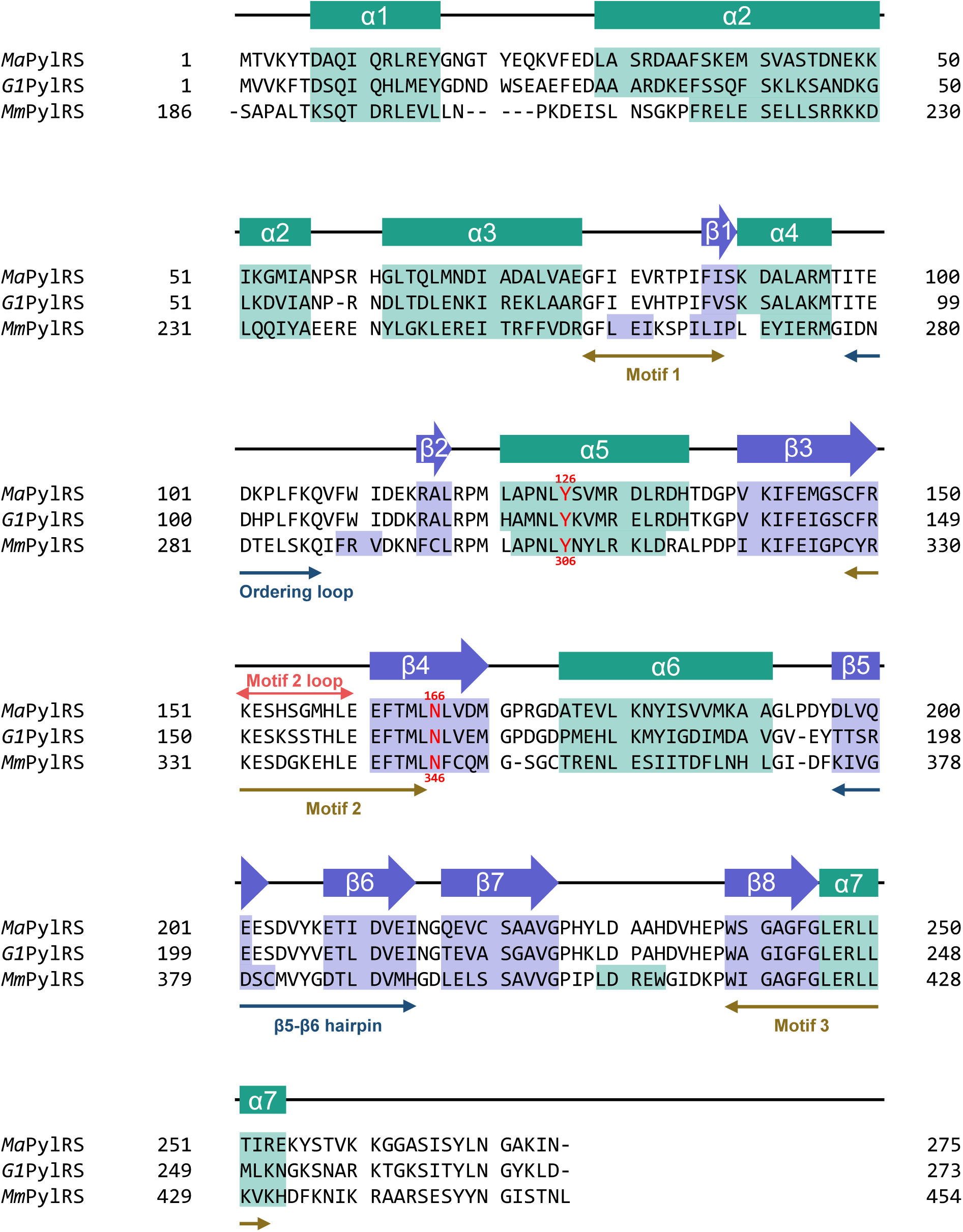
Sequence alignment and secondary structure annotations of *Ma*PylRS, *G1*PylRS, and *Mm*PylRS. The PylRS sequences were aligned using NCBI COBALT [PMID: 17332019]. The alignment includes *Ma*PylRS, *G1*PylRS and *Mm*PylRS. Secondary structure annotations are based on available crystal structures (PDB IDs: [6JP2], [8IFJ], and[2ZIM]) and previously reported annotations [PMID: 32182048, 37047230, and 17592110]. α-Helices and β-strands are shown above the alignment as green bars and purple arrows, respectively. Motifs 1–3, the ordering loop, the motif 2 loop, and the β5–β6 hairpin are indicated. Numbers flanking each sequence denote residue positions in the corresponding protein. Selected positions are additionally numbered above and below the alignment according to *Ma*PylRS and *Mm*PylRS numbering, respectively.

**Supplementary Figure 3.**
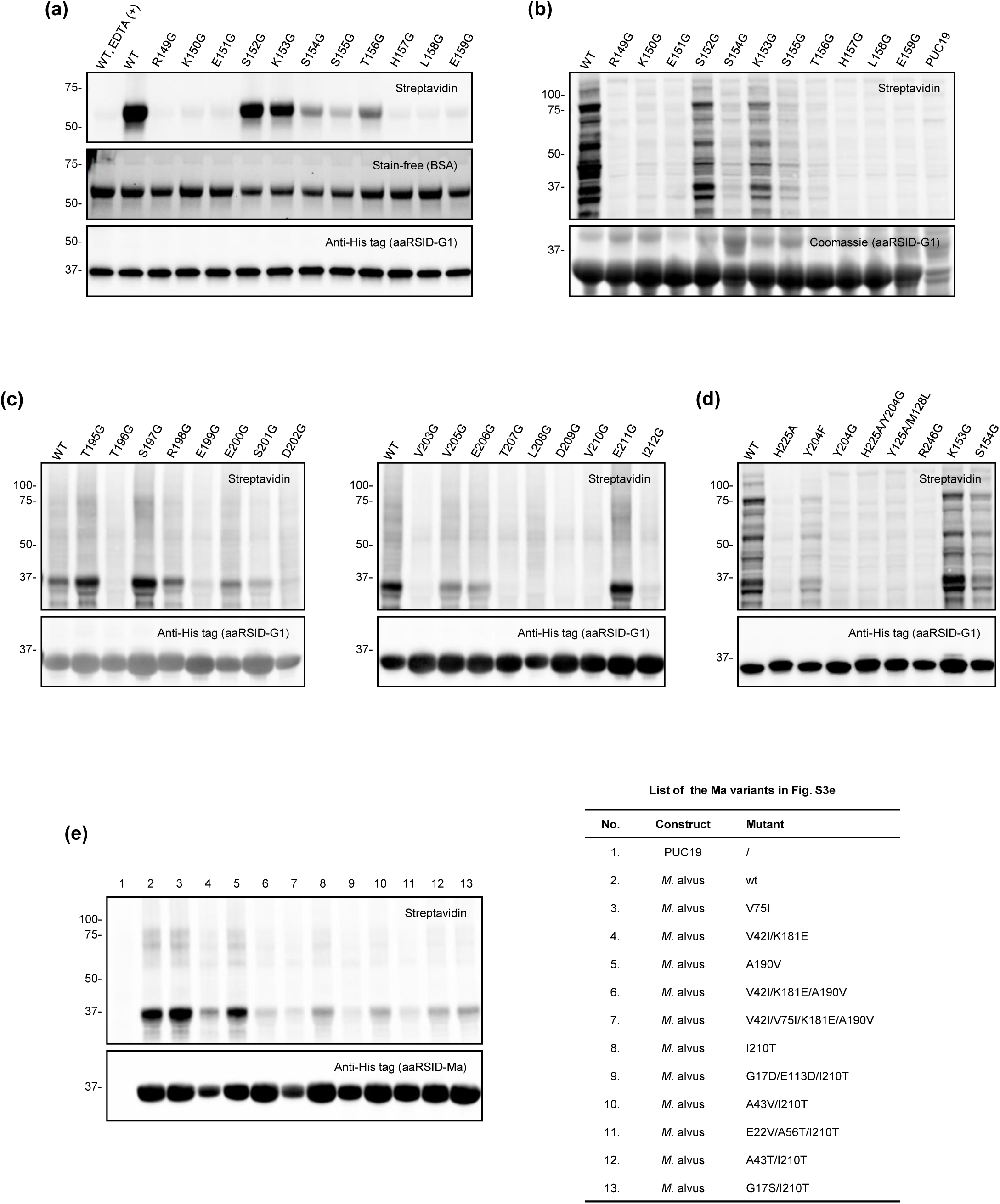
Screening of aaRSID variants for enhanced proximity labeling activity. (a, b) Screening of aaRSID-G1 glycine-substitution variants spanning the motif 2 loop for proximity labeling activity using an *in vitro* assay (a) and an *E. coli* assay (b). For the *in vitro* assay, 1 μg of each purified aaRSID-G1 variant was incubated with 150 μg BSA and 1 mM PrK, followed by CuAAC with biotin-azide and streptavidin blotting. An EDTA-containing reaction was included as a negative control. For the *E. coli* assay, *E. coli* BL21(DE3) cells expressing the indicated aaRSID variants were incubated with 1 mM PrK at 37 °C for 4 h, followed by protein extraction, CuAAC with biotin-azide, and streptavidin blotting. pUC19-transformed *E. coli* was included as a negative control. (c) Glycine-scanning mutagenesis of the β5–β6 hairpin of aaRSID-G1 using the *E. coli* assay. Variants spanning T195–I212 were screened for proximity labeling activity, with Y204G evaluated separately in (d). (d) Structure-guided follow-up analysis of selected aaRSID-G1 variants using the *E. coli* assay, including residues associated with β5–β6 hairpin conformational dynamics (Y204 and H225), substrate-binding pocket size (Y125A/M128L) (PMID: 37047230), and the conserved motif 3 residue R246 implicated in ATP binding (PMID: 18387634). (e) Screening of previously reported *Ma*PylRS variants evolved for PrK incorporation in GCE for their ability to support proximity labeling using the *E. coli* assay. Variant identities corresponding to lanes 1–13 are listed in the table on the right. pUC19-transformed *E. coli* was included as a negative control. (PMID: 35372501, PMID: 40413191)

**Supplementary Figure 4.**
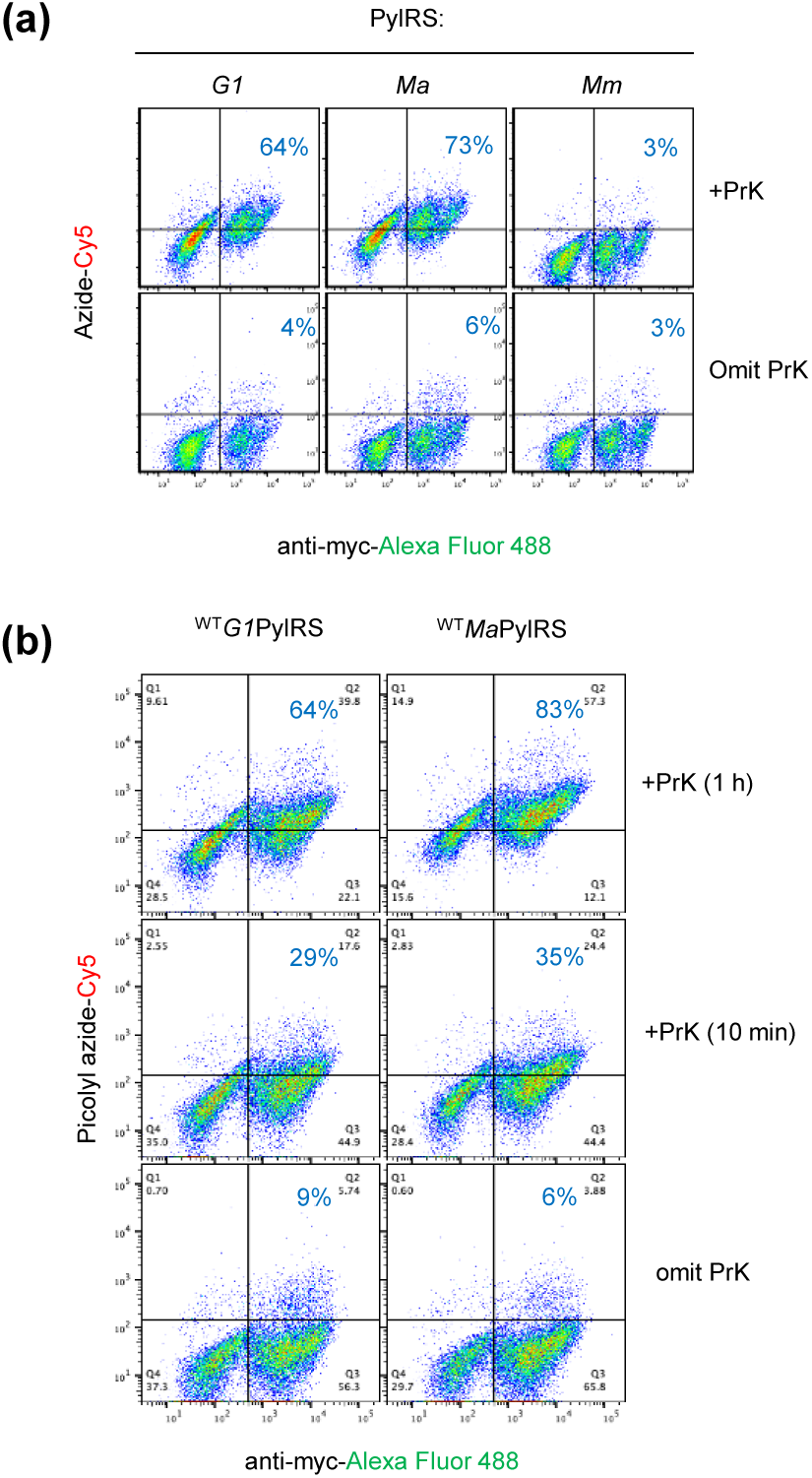
Surface display and labeling activity of PylRS homologs. (a) Cells displaying wild-type *G1*PylRS, *Ma*PylRS, or *Mm*PylRS were incubated with 1 mM PrK, 1 mM ATP, and 10 mM MgCl₂ for 1 h. Reactions performed in the absence of PrK served as negative controls. PrK-modified cells were detected by CuAAC with picolyl-azide-Cy5, while PylRS surface expression was detected using an anti-Myc antibody. Samples were analyzed by FACS, and labeling activity was quantified as the percentage of labeling-positive cells (Q2) among total PylRS-expressing cells (Q2 + Q4). (b) Cells were incubated with 1 mM PrK, 1 mM ATP, and 10 mM MgCl₂ for 10 min and 1 h. Reactions performed in the absence of PrK served as negative controls. PrK-modified cells were detected by CuAAC with picolyl-azide-Cy5, while PylRS surface expression was detected using an anti-myc antibody. Samples were analyzed by FACS, and labeling activity was quantified as the percentage of labeling-positive cells (Q_2_) among total PylRS-expressing cells (Q_2_ + Q_4_).

**Supplementary Figure 5.**
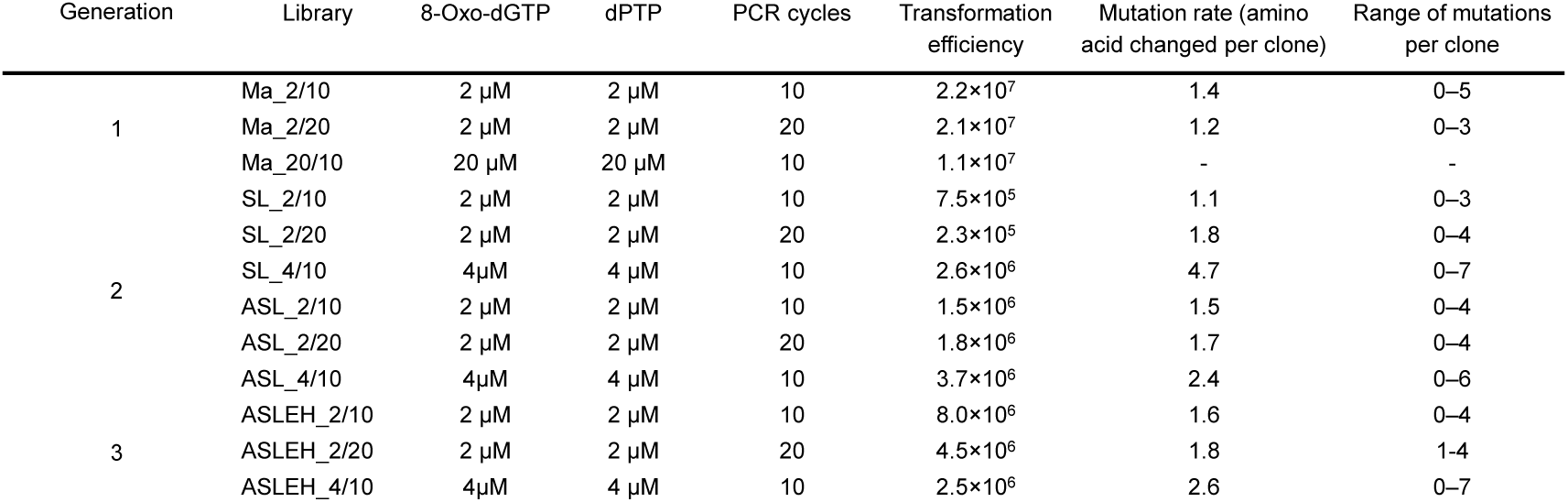
Summary of error-prone PCR conditions used to generate three generations of *Ma*PylRS libraries and the resulting library parameters. Transformation efficiency was determined by plating serial dilutions of each library on selective agar. Mutation frequencies and ranges were determined by sequencing 20 randomly selected clones from each library.

**Supplementary Figure 6.**
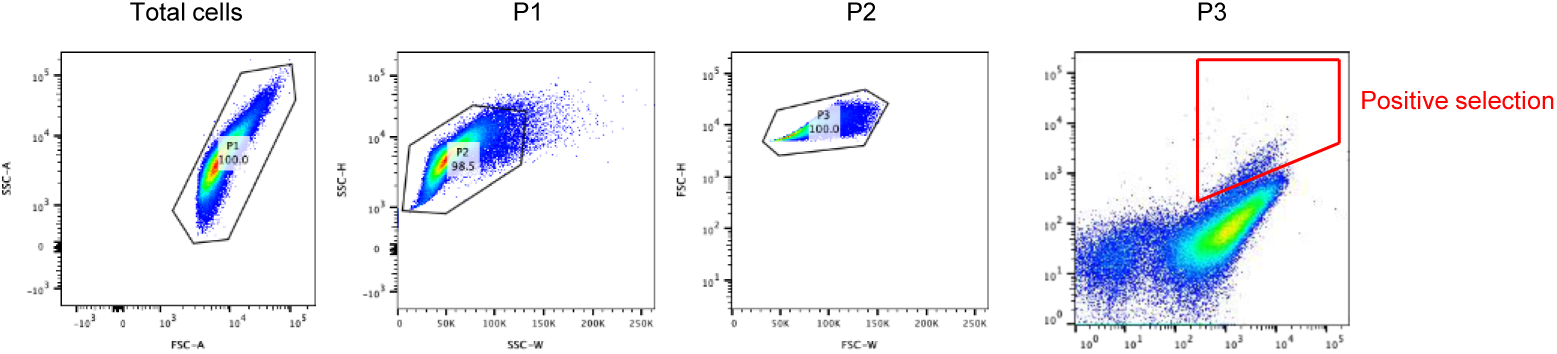
Gating strategies for isolating single cells and positive selection. Total cells were gated around the clustered population on a forward-scatter area (FSC-A) versus side-scatter area (SSC-A) to yield population 1 (P1). P1 was then resolved on a side-scatter width (SSC-W) versus side-scatter height (SSC-H) plot to isolate population 2 (P2). P2 was plotted on a forward-scatter width (FSC-W) versus forward-scatter height (FSC-H) plot to yield the single-cell population 3 (P3). P3 single cell population were then plotted using the X-axis for protein expression (anti-Myc) and the Y-axis for labeling activity (streptavidin conjugate). A trapezoidal gate was drawn to collect the cells exhibiting high streptavidin/anti-myc ratios.

**Supplementary Figure 7.**
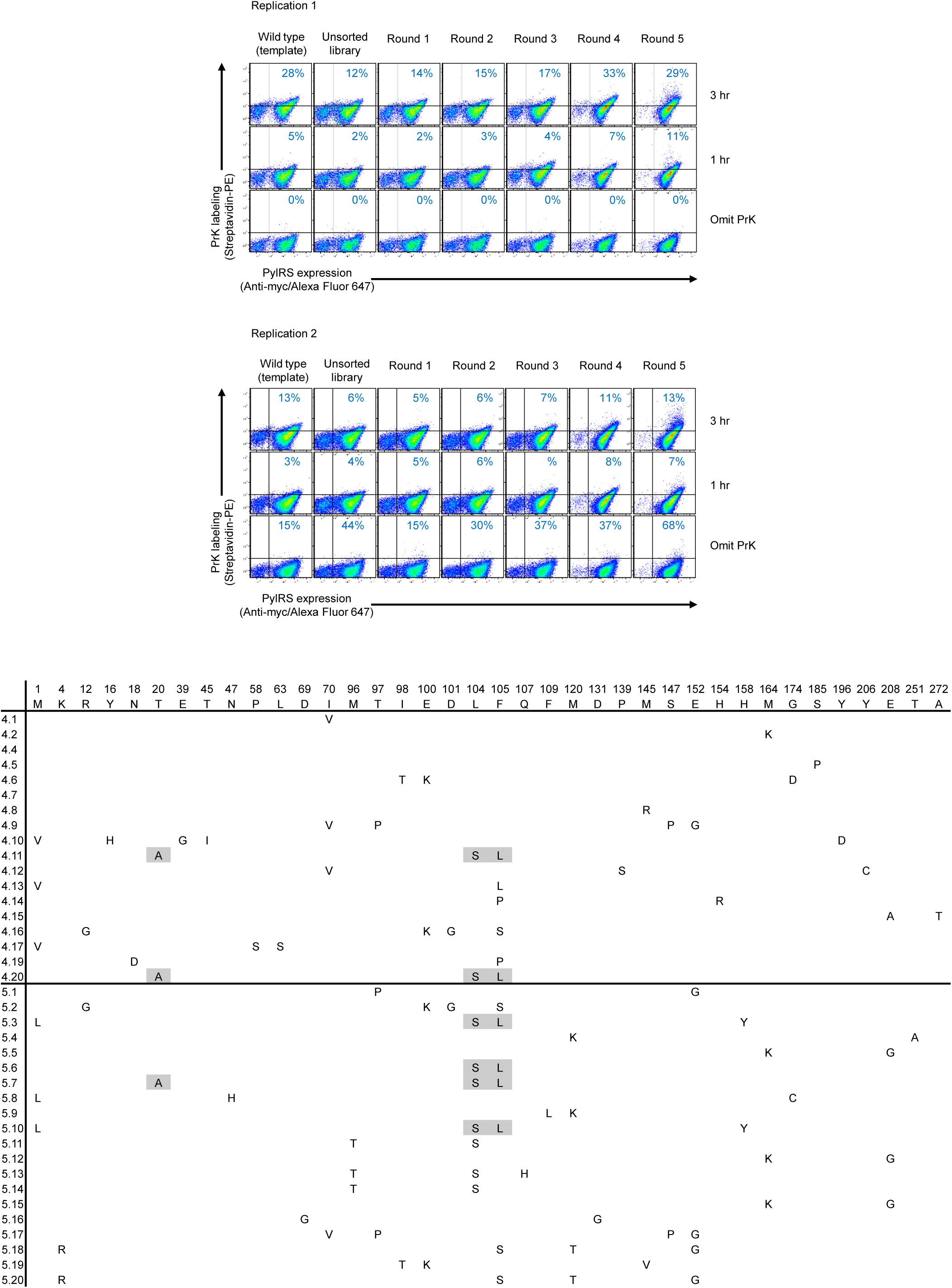
Generation 1 directed evolution of aaRSID using yeast display. **(a)** Yeast libraries from each selection round were labeled under identical conditions (1 mM PrK, 1 mM ATP, and 10 mM MgCl₂). The labeling duration was progressively reduced over five rounds of selection (6, 3, 3, 3, and 1 h, respectively). Following labeling, cells were subjected to CuAAC with azide-biotin and stained with anti-Myc antibody and streptavidin– phycoerythrin prior to FACS analysis. Two independent replicates were performed, with 50,000 cells (top) and 100,000 cells (bottom). **(b)** Sequence analysis of enriched variants after rounds 4 and 5 of yeast display selection. Plasmids were extracted from pooled yeast populations and transformed into *E. coli*. Twenty individual colonies from each round were analyzed by Nanopore whole-plasmid sequencing. Amino acid substitutions identified in selected variants relative to wild-type MaPylRS are summarized in the table above.

**Supplementary Figure 8.**
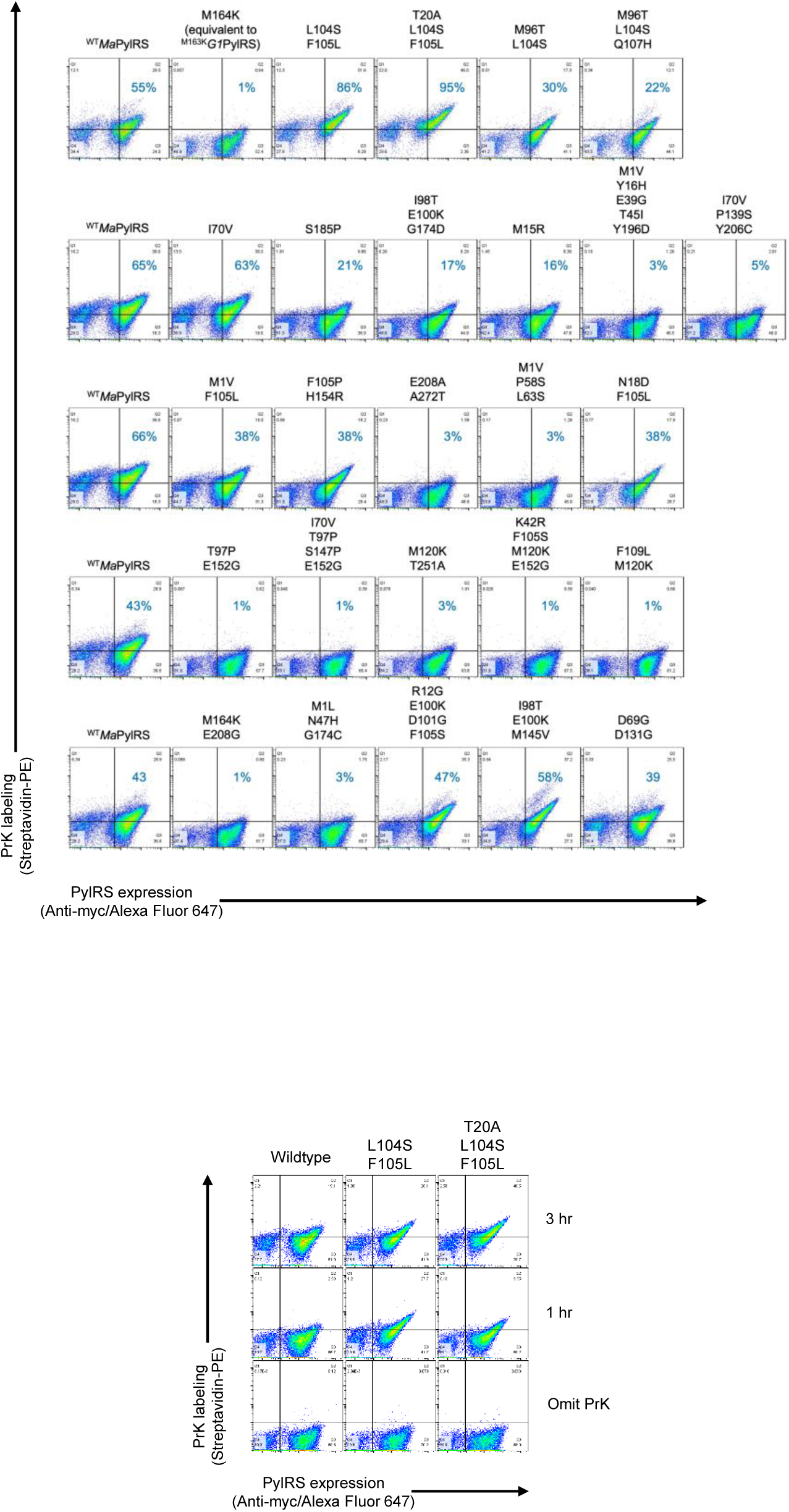
Individual clone characterization. Individual clones identified from round 4 and 5 were expressed on yeast surface and compared for labeling activity side-by-side with the wild type.

**Supplementary Figure 9.**
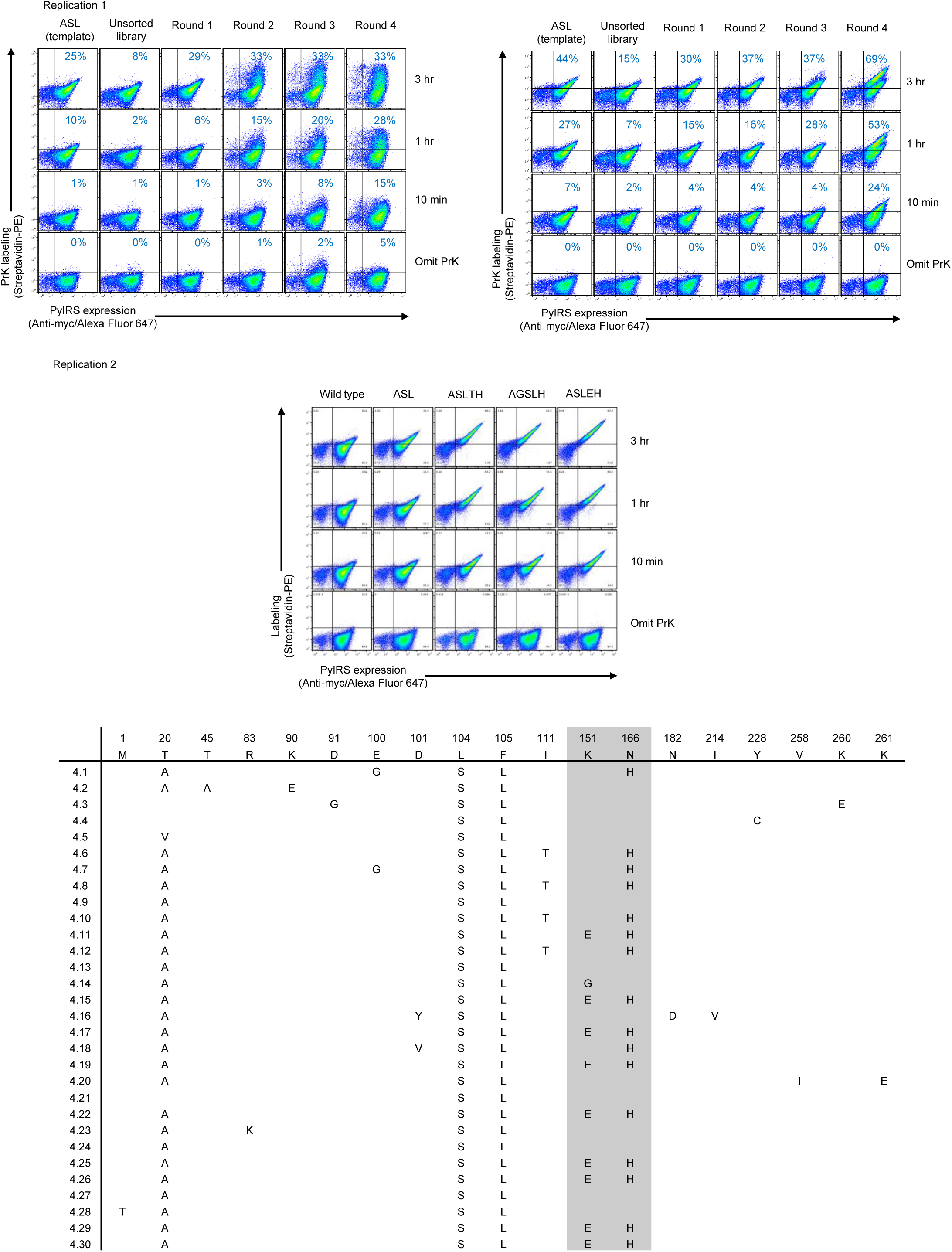
Generation 2 for directed evolution of aaRSID using yeast display. Pooled yeast library each round was labeled using the same condition: 1 mM PrK, 1 mM ATP and 10 mM MgCl2 in 3% BSA-PBS. Five rounds of selection were performed with decreased labeling time: 6, 3, 3, 3 and 1 h. Labeled cells were later detected via CuAAC with azide-biotin followed by anti-myc antibody and streptavidin-phycoerythrin labeling before FACS analysis. This experiment was performed with 2 replicates with 50,000 cells (above) and 100,000 cells (below) analyzed.

**Supplementary Figure 10.**
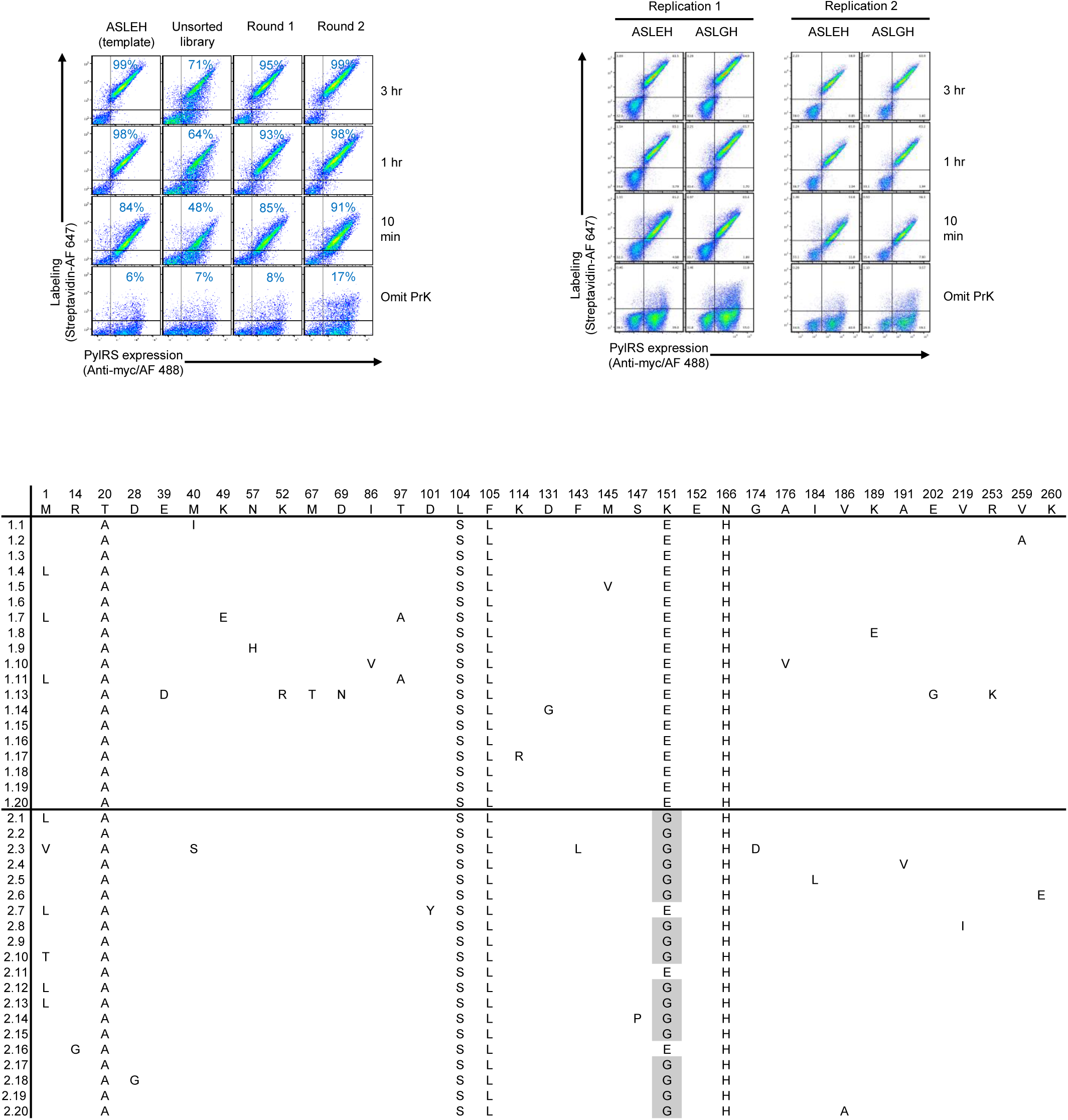
Generation 3 for directed evolution of aaRSID using yeast display. Pooled yeast library each round was labeled using the same condition: 1 mM PrK, 1 mM ATP and 10 mM MgCl2 in 3% BSA-PBS. Five rounds of selection were performed with decreased labeling time: 6, 3, 3, 3 and 1 h. Labeled cells were later detected via CuAAC with azide-biotin followed by anti-myc antibody and streptavidin-phycoerytrin labeling before FACS analysis. This experiment was performed with 2 replicates with 50,000 cells (above) and 100,000 cells (below) analyzed.

**Supplementary Figure 11.**
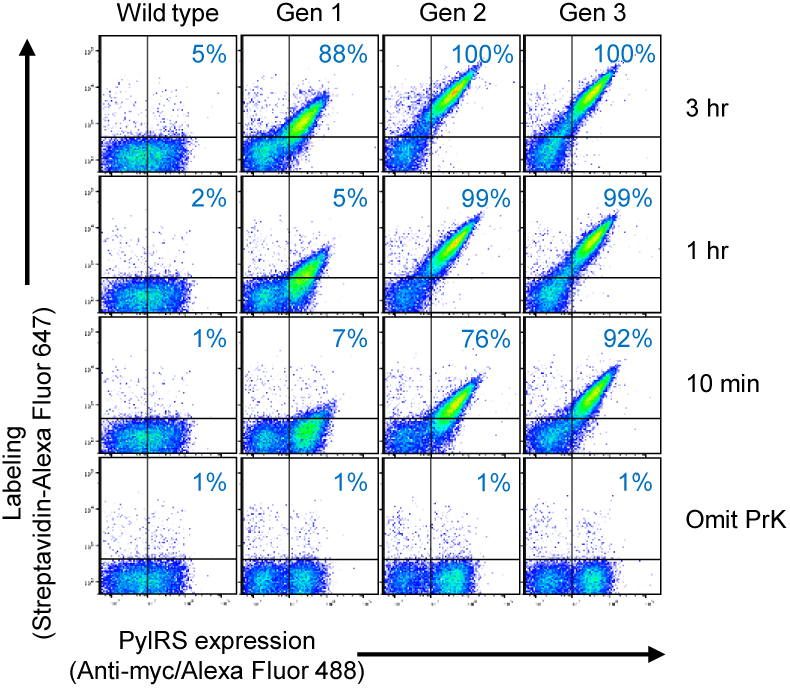
Comparison of the winning clones from each directed evolution generation. Cells were labeled with 1 mM PrK, 1 mM ATP, and 10 mM MgCl_2_ for the designated time points (10 min, 1 h and 3 h). Negative control (omit PrK) was performed as without PrK added for 3 h. This experiment was performed once.

**Supplementary Figure 12.**
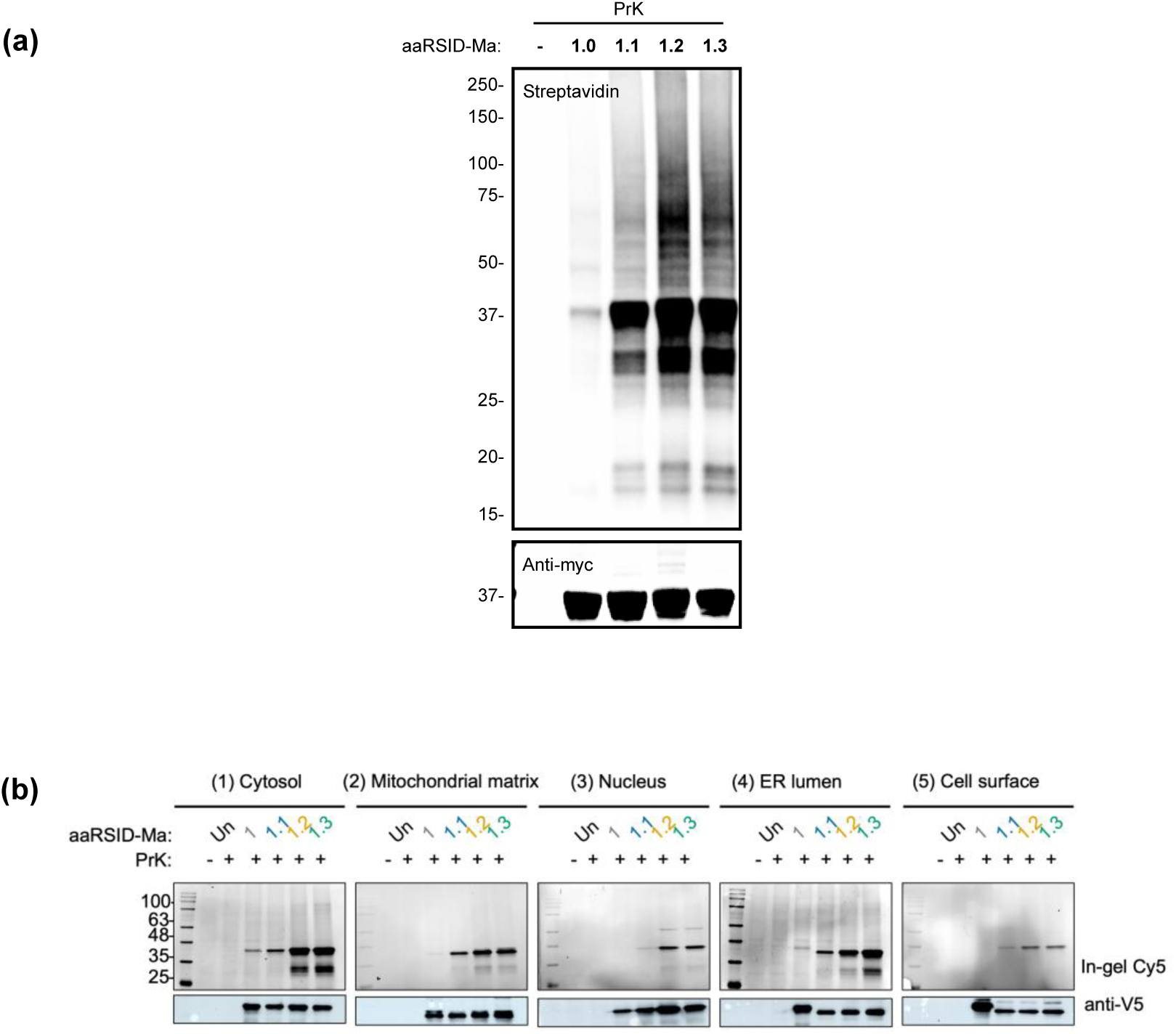
Comparison of labeling activity of evolved aaRSID-Ma variants across subcellular compartments in HEK293T cells. (a) Streptavidin blotting showing PrK labeling in cells expressing the indicated aaRSID-Ma1.X-NES variants. Following labeling, proteins were conjugated to biotin-azide via CuAAC. (b) Cells expressing aaRSID-Ma in different subcellular localizations were labeled with 1 mM PrK for 6 h. Whole-cell lysates were stained with azide-Cy and analyzed by in-gel Cy5 fluorescence, while enzyme expression was measured by anti-V5 blotting.

**Supplemental Figure 13.**
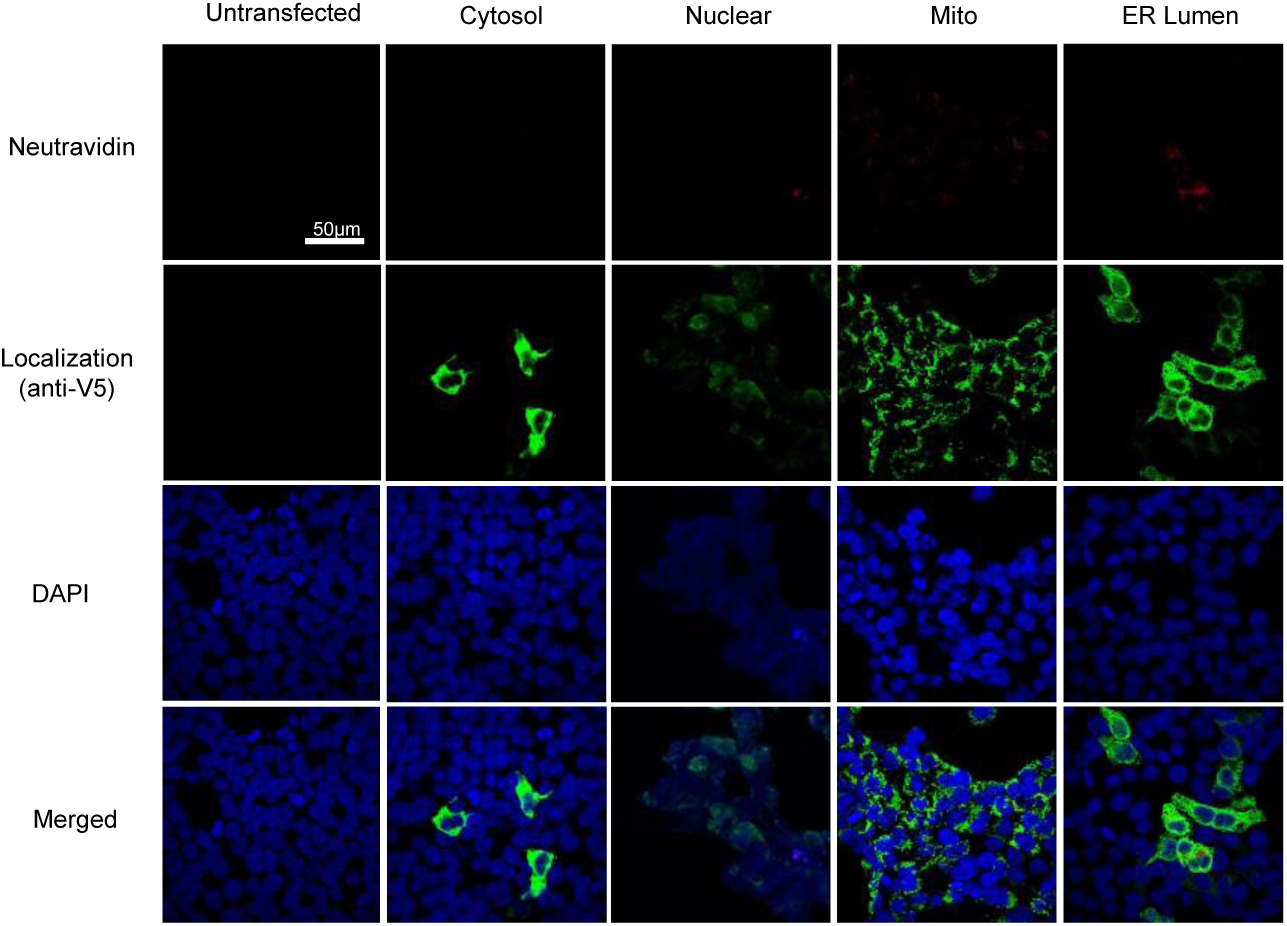
Characterization of wild-type PylRS proximity labeling via confocal fluorescence imaging. Transfected cells were labelled by PylRS and PrK for 24hr. PrK labeled proteins were detected from fixed cells using a biotin azide probe via copper-catalyzed azide-alkyne cycloaddition (CuAAC). Labeled proteins and aaRSID expression were detected using streptavidin Dylight 594 and AlexaFluor 488 anti-V5, respectively. Labeling was normalized to the cells depicted in Figure 4d.

**Supplemental Figure 14.**
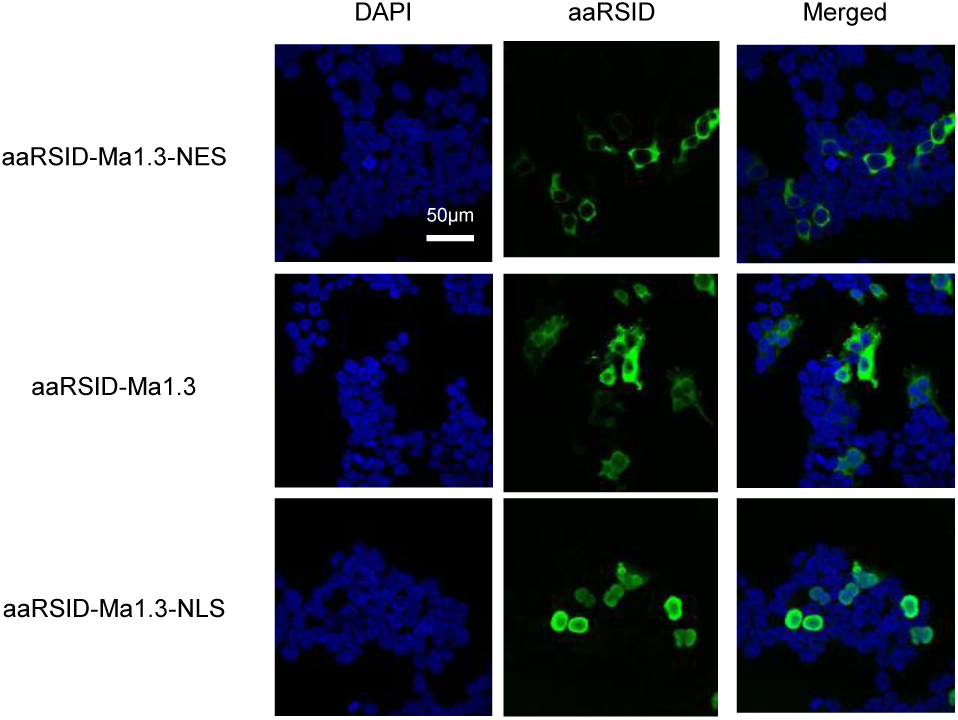
Localization test of aaRSID mutant. HEK293T cells were transiently transfected with the aaRSID-Ma1.3 mutant without a localization sequence (middle). Cells were transfected with the cytoplasmic (top) and nuclear (bottom)-localized constructs as a control. aaRSID expression was detected with AlexaFluor488, while DAPI was used to stain the nucleus. Images are normalized to each other.

**Supplementary Figure 15.**
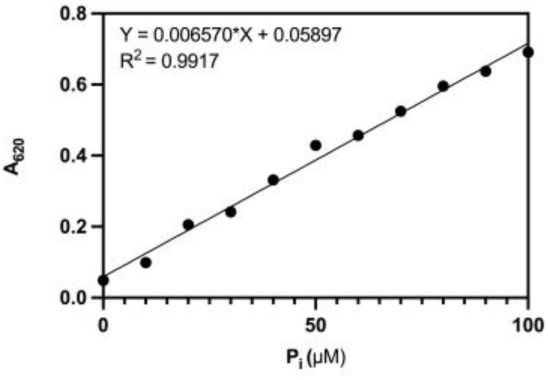
Calibration curve of inorganic phosphate. A phosphate standard solution was prepared by dissolving KH_2_PO_4_ in water and diluted to 0-100 µM. Each standard dilution was then reacted with malachite green/molybdate solution for 30 min. The absorbance was determined at 620 nm using a microplate reader.

**Supplementary Figure 16.**
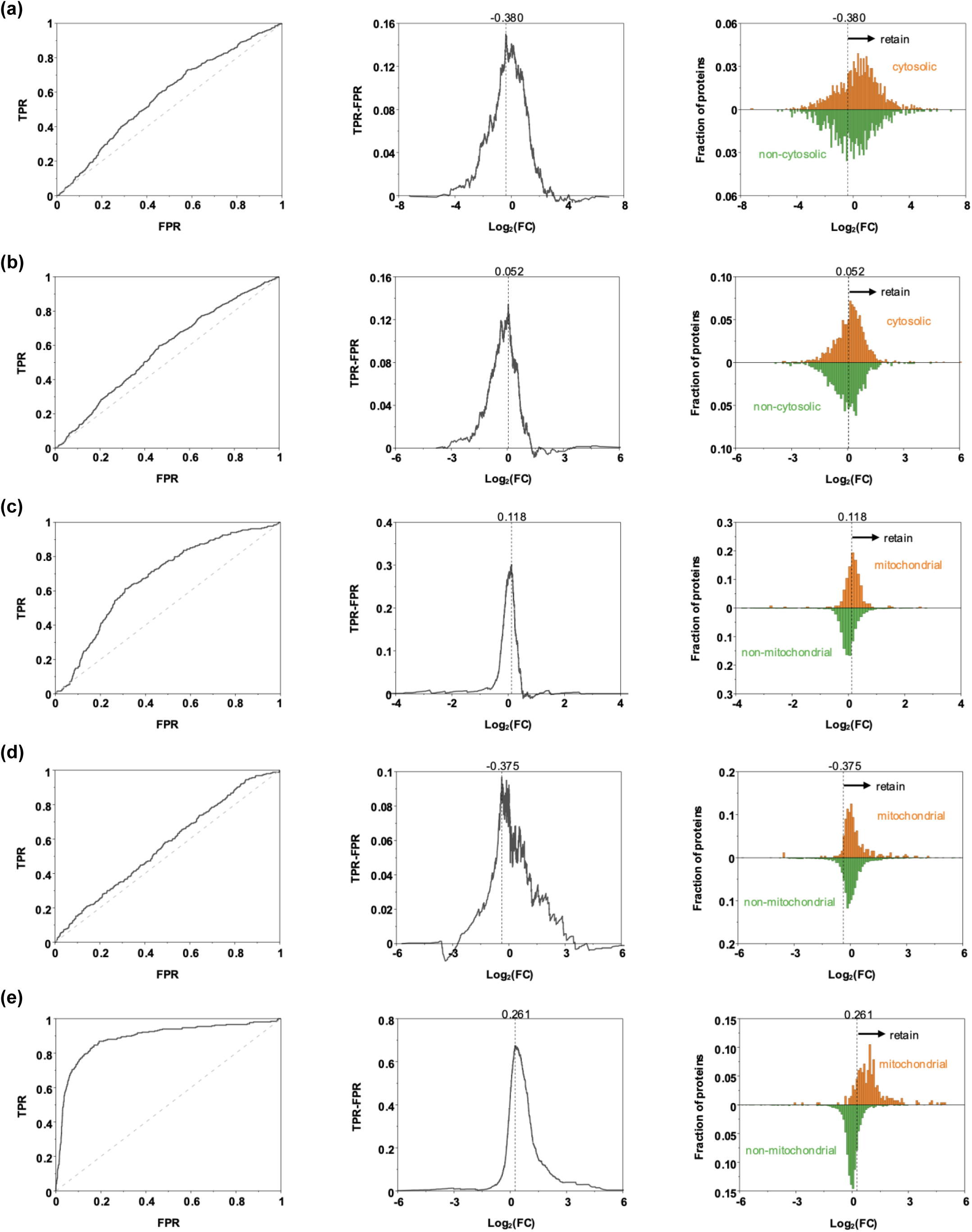
(a, b) ROC-based filtering of aaRSID-NES proteomic data obtained using the biotin-azide/streptavidin (a) and FAM-azide/anti-FAM (b) workflows, related to Figure 5b, d. (c∼e) ROC-based filtering of proteomic data obtained using aaRSID-Mito (c), TurboID-Mito (d), and APEX2-Mito (e), related to Figure 5c, e. Left, ROC curves generated using annotated positive and negative reference protein sets. Middle, true-positive rate minus false-positive rate (TPR−FPR) plotted against log₂(fold change), with the maximum used to define the filtering threshold. Right, distributions of log₂(fold change) for true positive (orange, plotted upward) and false positive (green, plotted downward) reference protein sets. Vertical dashed lines indicate the selected thresholds; arrows indicate the retained range.

